# Mechanism of an alternative splicing switch mediated by cell-specific and general splicing regulators

**DOI:** 10.1101/2023.01.23.525191

**Authors:** Yi Yang, Giselle C Lee, Erick Nakagaki-Silva, Yuling Huang, Matthew Peacey, Ruth Partridge, Clare Gooding, Christopher WJ Smith

## Abstract

Alternative pre-mRNA splicing is regulated by RNA binding proteins (RBPs) that activate or repress regulated splice sites. Repressive RBPs bind stably to target RNAs via multivalent interactions, which can be achieved by both homo-oligomerization and by interactions with other RBPs mediated by intrinsically disordered regions (IDRs). Cell-specific splicing decisions commonly involve the action of widely expressed RBPs that can bind around target exons, but without effect in the absence of a key cell-specific regulator. To address how cell-specific regulators collaborate with constitutive RBPs in alternative splicing regulation we used the smooth-muscle specific regulator RBPMS. Recombinant RBPMS is sufficient to switch cell specific alternative splicing of *Tpm1* exon 3 in cell free assays by remodelling ribonucleprotein complexes and preventing assembly of ATP-dependent splicing complexes. This activity depends upon its C-terminal IDR, which facilitates dynamic higher-order self-assembly, cooperative binding to multivalent RNA, and interactions with other splicing co-regulators, including MBNL1 and RBFOX2. Our data show how a cell-specific RBP can co-opt more widely expressed regulatory RBPs to facilitate cooperative assembly of stable cell-specific regulatory complexes.

## Introduction

Alternative pre-mRNA splicing (AS) is a widespread phenomenon in eukaryotes that allows multiple transcripts to be generated from individual genes, often leading to the production of functionally distinct protein isoforms with profound effects on cell and organismal phenotype (Kalsotra & Cooper, 2011, Wright et al., 2022). Genome-wide studies demonstrate that most AS events (ASEs) are mediated by the combinatorial and tissue-specific binding of multiple RNA binding proteins (RBPs) (Huelga et al., 2012, Pandit et al., 2013, Warzecha et al., 2009). The human genome encodes at least 1500 RBPs (Dominguez et al., 2018, Gerstberger et al., 2014), many of which comprise one or more RNA binding domains (RBDs) along with a variety of intrinsically disordered regions (IDR) (Corley et al., 2020). While much focus has been placed on the role of structurally ordered RBDs in the recognition of specific RNA motifs (Dominguez et al., 2018), recent studies have also begun to unveil the biological significance of the IDRs (Lin et al., 2015, Protter et al., 2018).

Most splicing regulatory RBPs have preferred binding motifs, which act as splicing enhancers or silencers depending on the activity of the cognate RBP (Fu & Ares, 2014). RBPs work either synergistically or antagonistically to modulate spliceosome assembly at regulated splice sites, resulting in either exon activation or repression (Dvinge, 2018, Jangi & Sharp, 2014, Ule & Blencowe, 2019). Tissue-specific “splicing codes” comprise different combinations of enhancer and silencer motifs along with other transcript features (Barash et al., 2010, Fu & Ares, 2014). Most features of cell-specific splicing codes are not unique to that cell type, reflecting the roles of widely expressed RNA binding proteins in cell specific splicing decisions (Barash et al., 2010). Nevertheless, some splicing regulators show more restricted expression and may act as master regulators of splicing programs (Jangi & Sharp, 2014, Raj et al., 2014). The outcome of splicing decisions can be viewed as resulting from a competition between activating and repressive inputs, while switching between AS patterns can result from modulation of either or both sets of inputs. For example, many neuron-specific exons are included as a result of reduced repression by PTBP1 combined with increased activation by RBFOX or SRRM3 proteins (Barash et al., 2010, Castle et al., 2008, Raj et al., 2014, Wang et al., 2008).

Splicing activators of the SR protein family can act by binding to exon splicing enhancers (ESEs) via their RRM domains, while their serine-arginine (SR) rich IDRs either recruit core splicing factors to splice sites or stabilize interactions within splicing complexes (discussed in (Jobbins et al., 2018)). Increased numbers of ESEs additively enhance splicing efficiency, but this arises from increased probability of initial weak binding to ESEs (Jobbins et al., 2018) or increased probability of interaction of ESE bound SR proteins with core splicing factors (Hertel & Maniatis, 1998) rather than co-operation between SR proteins bound to different ESEs. Splicing repressors broadly act in one of two ways. Their binding can directly occlude splice sites, ESEs, or whole exons, blocking the binding of activating or core splicing factors (Fu & Ares, 2014, Ule & Blencowe, 2019). Alternatively, repressors can interact with RNA bound core splicing factors leading to dead-end splicing complexes (Chiou et al., 2013, Sharma et al., 2008, Sharma et al., 2012). Exemplified by the hnRNP family, repressors have one or more RBDs and typically interact in a multivalent manner with target RNAs containing multiple cognate binding motifs. Multivalency can arise via multiple RBDs within a single protein or via oligomerization mediated by intrinsically disordered regions (IDRs) (Ule & Blencowe, 2019). The IDRs have a propensity for mediating both homotypic and heterotypic protein-protein interactions, including higher-order oligomerization and biomolecular condensate formation, and have been shown to be functionally important in a range of splicing regulators such as RBFOX2 (Damianov et al., 2016, Sun et al., 2012, Ying et al., 2017), hnRNPH1 (Kim & Kwon, 2021), hnRNPA and hnRNPD (Gueroussov et al., 2017). It has been proposed that some RBPs might act by promoting local “binding region condensates” on target transcripts (Hallegger et al., 2021).

Detailed mechanistic understanding of the action of splicing regulatory RBPs can be gained from cell free *in vitro* investigations. For example, biochemical investigations of the *CSRC* N1 exon have provided a detailed picture of how the archetypal repressor PTBP1 leads to exon skipping via cooperative binding to motifs flanking the exon (Chou et al., 2000), leading to hyperstabilized non-productive U1 snRNP binding at the N1 5’ss (Sharma et al., 2008, Sharma et al., 2012). Here, PTBP1 acts widely as a splicing repressor and its reduced expression in neurons leads to N1 exon inclusion. *In vitro* analyses of the action of cell-specific regulators are lacking, possibly due to challenges associated with expression and purification of active full-length proteins with extensive IDRs. We recently found that the 22 kDa RNA binding protein RBPMS is sufficient to activate a splicing program associated with differentiated contractile vascular smooth muscle cells (VSMCs) (Nakagaki-Silva et al., 2019). Among the AS events regulated by RBPMS was the switch between *Tpm1* mutually exclusive exons 2 and 3, an event that has been extensively investigated using *in vitro, in cellulo* and *in vivo* approaches (Gooding & Smith, 2008). *Tpm1* exon 3 inclusion results from its dominant 5′ and 3′ splice site (5′ss and 3′ss) elements, which outcompete the weaker exon 2 splice sites, except in differentiated SMCs where exon 3 is repressed (Gooding et al., 1994, Mullen et al., 1991). MBNL and PTBP1 proteins apply a constitutive repressive influence on exon 3 by binding to flanking negative regulatory sequences (Cherny et al., 2010, Ellis et al., 2004, Gooding et al., 2013, Gooding et al., 1998). However, both proteins are widely expressed (Llorian et al., 2016) and despite the binding of up to 6 PTBP1 and 3-8 MBNL1 proteins bind around *Tpm1* exon 3 it is efficiently included in HeLa nuclear extract splicing reactions (Cherny et al., 2010, Gooding et al., 2013, Gooding et al., 1998). Since RBPMS over-expression is sufficient to switch *Tpm1* splicing in cell lines such as HEK293T (Nakagaki-Silva et al., 2019) we hypothesised that recombinant RBPMS might be able to confer tissue-specific splicing of *Tpm1* in cell-free assays. RBPMS has a single RNA recognition motif (RRM) that mediates both homodimerization (Sagnol et al., 2014, Teplova et al., 2015) and binding to closely spaced pairs of CAC motifs (Farazi et al., 2014, Soufari & Mackereth, 2017), a 14 aa N-terminal tail, and an ∼80 aa proline-rich C-terminal IDR (Fig. 1A). The IDR is important for some functions (Aguero et al., 2016, Gerber et al., 2002, Kaufman et al., 2018) and can contribute to RNA binding (Farazi et al., 2014), but the biophysical basis of its activity is unclear.

**Figure 1.**
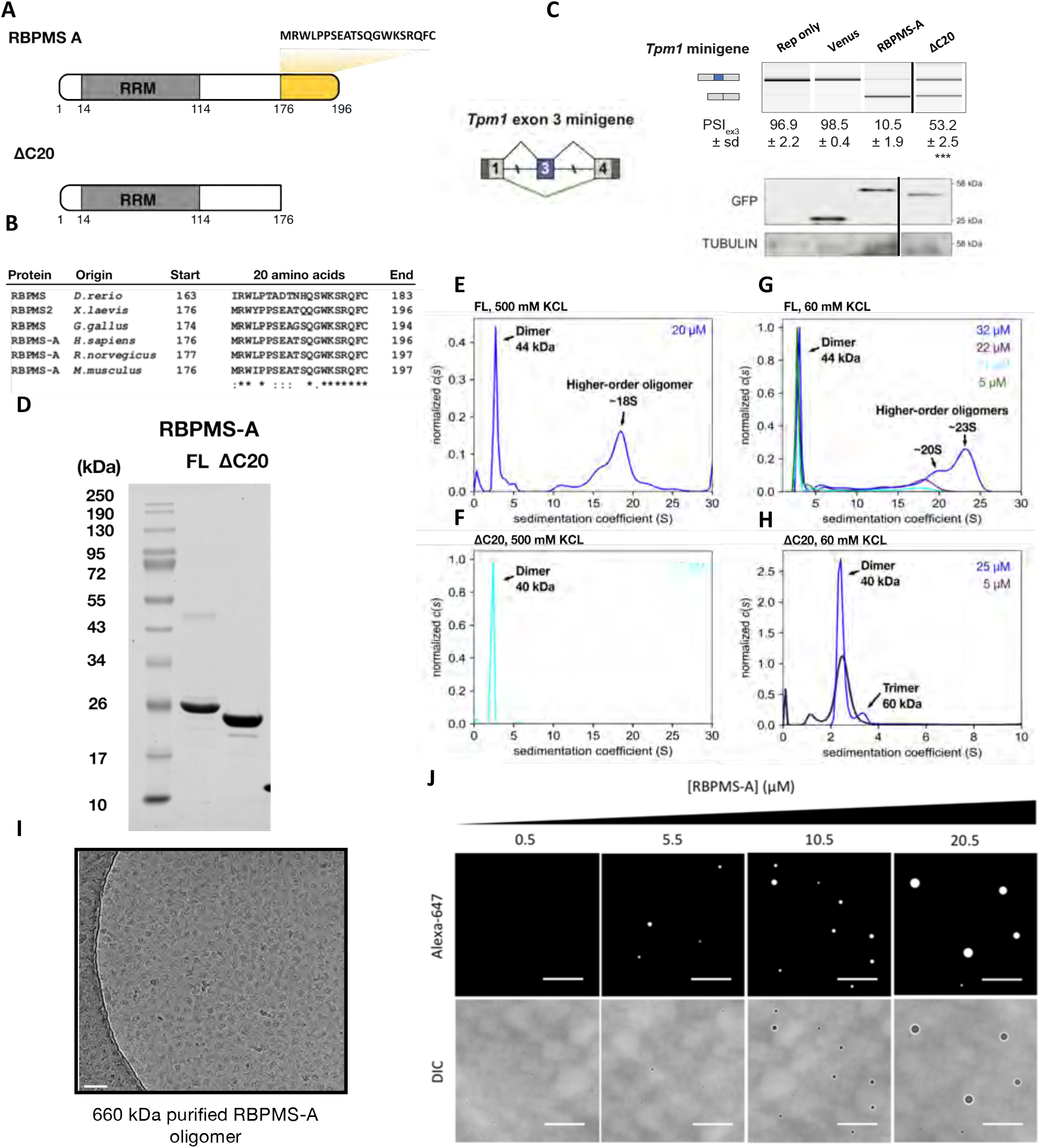
*In vitro* characterization of recombinant RBPMS-A. (A) Alternative exon encodes 20 amino acids RBPMS-A isoform specific tail (yellow). αC20, an experimental construct lacking the C-terminal tail. (B) Homology of C-terminal 20 amino acids of RBPMS orthologs from vertebrates. Alignment was generated using Clustal Omega with sequences : NP_001008710.1 ; NP_001036139.1 ; XP_015132108.2 ; NP_001081864.1 ; XP_006253302.1 and ; XP_003199078.1. The degree of conservation is indicated at the bottom of the sequences where asterisk indicates fully conserved residues, colon indicates residues of strongly similar properties, and period indicates residues of weakly similar properties.(C) Tpm1 exon 3 minigene reporter co-transfected with Venus tagged RBPMS in HEK293 cells. Schematic of the minigene is shown on the left. RT-PCRs for the splicing patterns. PSI values are shown as mean ±sd from n=3. Differentiated splicing is shown in green, proliferative exon is in blue. Statistical significance from Student’s t-test is shown as *** = p < 0.001.(D) SDS-PAGE analysis of purified recombinant proteins. RBPMS-A full-length (FL) and C-terminal 20 amino acids truncated (αC20) (E-H) Sedimentation coefficient distribution plot, *c(s)* of AUC analysis using FL or αC20 RBPMS at 500 mM or 60 mM KCL concentration, pH7.9. Protein concentrations are color-coded. Data shown in E and F are normalized to the area under curve using GUSSI, while for G and H data are normalized to the maximum value of the dataset. (I) Cryo-electron microscopy image of glutaraldehyde cross-linked and purified RBPMS-A high-order oligomer of 660 kDa. Scale bar, 50 nm. (J) Fluorescence and differential interference contrast micrographs of His6-RBPMS-A LLPS droplets at 90 mM KCL. Fluorophore-conjugated RBPMS-AAlexa-647 is added to 0.5 μM to give the final concentration shown. Images are representative of 5-10 acquired at each concentration. Scale bars are 25 μm.

Here, we show that recombinant RBPMS confers cell-specific AS of *Tpm1* exon 3 *in vitro* by remodelling the ribonucleoprotein (RNP) complexes that assemble around the exon, thereby preventing the formation of ATP-dependent splicing complexes. The IDR is essential for RBPMS splicing regulatory function and for its ability to bind to *Tpm1* RNA in nuclear extract. It mediates higher-order oligomerization extending to liquid-liquid phase separation, cooperative binding to the multivalent *Tpm1* RNA, as well as interaction with other widely expressed splicing regulators such as MBNL1 and RBFOX2. In particular, the interaction with MBNL1 helps to recruit RBPMS to *Tpm1* RNA in the competitive context of nuclear extract, while RBFOX2 and other proteins are recruited by RBPMS. Notably both MBNL1 and RBFOX2 co-regulate not only of *Tpm1* but also other VSMC regulated events. Our results provide an important proof of principle for how a cell-specific splicing regulator can interact functionally and physically with more widely expressed regulators to direct their activity towards a coregulated set of AS events.

## Results

### RBPMS assembles into higher-order oligomeric structures via its C-terminus

Vertebrate RBPMS and RBPMS2 sequences show high conservation of the C-terminal 20 amino acids corresponding to the major RBPMS isoform (Fig. 1A, with complete conservation of aromatic and basic residues (Fig. 1B). Deletion of this region in transfected RBPMS (αC20) led to a significant loss in splicing repressor activity (Fig. 1C), demonstrating that the RRM alone is insufficient for RBPMS splicing regulatory function, despite being sufficient for dimerization and sequence-specific binding (Sagnol et al., 2014, Teplova et al., 2015). For *in vitro* analyses we prepared recombinant untagged full length and ΔC20 rat RBPMS (Fig. 1D). Around 60% of full length (FL) RBPMS eluted as a dimer during size exclusion chromatography (SEC), consistent with previous reports (Fig. S1A) (Sagnol et al., 2014, Teplova et al., 2015). The remaining 40% eluted as a broad peak before the 443 kDa marker approaching the void volume, indicative of heterogeneous higher-order assemblies. In contrast, 96% of the ΔC20 mutant eluted as a single peak corresponding to the dimeric state, suggesting that the C-terminal region is required for RBPMS higher-order oligomerization. Both proteins migrated corresponding to their monomeric molecular weight on SDS PAGE (Fig. 1D).

To examine RBPMS higher-order structures in more detail, we carried out sedimentation velocity analytical ultracentrifugation (AUC). At high ionic strength (500 mM KCl), FL RBPMS displayed polydispersity and existed as a series of dimers and higher-order oligomers (Fig. 1E). The largest oligomeric species sedimented at 18S, corresponding to approximately ∼460 kDa (∼21 monomers). In contrast, ΔC20 sedimented homogeneously as a dimeric protein (Fig. 1F), consistent with its SEC profile (Fig S1B). At low salt conditions used for *in vitro* splicing assays (60 mM KCl), FL RBPMS oligomerized in a concentration-dependent manner. Oligomers as large as 23S (∼600 kDa) were observed at 32 μM (Fig. 1G). In contrast, the ΔC20 mutant was mainly dimeric, with a small amount of trimers forming at 25 μM (Fig. 1H). Using glycerol gradients supplemented with a glutaraldehyde crosslinker (GraFix), we found that the C-terminal tail is required for the formation of higher-order oligomers above 250 kDa, but not trimeric and tetrameric species, which were more prominent with ΔC20 (Fig. S3). FL RBPMS and ΔC20 also exhibit different friction ratios that reflect potential shape differences; FL RBPMS and ΔC20 dimers have friction ratios over 1.4 (Fig. S2E-H), whereas heterogeneous RBPMS oligomers have friction ratios of 1.0 to 1.15 (Fig. S2A-D). This indicates that RBPMS dimers are more elongated in conformation, whereas RBPMS higher-order structures are likely more spherical. We confirmed the spherical shape of RBPMS higher-order structures by subjecting chemically cross-linked oligomers of ∼660 kDa to cryoEM (Fig. 1I, Fig. S3). However, 2-dimensional projections of RBPMS oligomers were unamenable to further classification, indicating a high degree of structural heterogeneity. Consistent with this, His_6_-tagged and tag-free RBPMS were observed to form droplets in vitro (Fig. 1J and S4B). The phase separation of tag-free RBPMS required the presence of the molecular crowding reagent PVA (Fig S4B). To examine the nature of phase-separated RBPMS droplets, His_6_-tagged RBPMS was fluorescently labeled with Alexa Fluor 647 (Fig S4A), spiked into unlabeled RBPMS, and monitored by fluorescence microscopy. Pre-formed RBPMS droplets acquired fluorescence after mixing and showed protein concentration-dependent changes in volume, suggesting that they are liquid-like and dynamic. Taken together, our biophysical analyses show that RBPMS exists as a heterogeneous mixture of concentration-dependent higher-order oligomers in addition to dimers, and that the C-terminal 20 amino acids that are important for activity *in vivo* are also essential for higher order oligomerization.

### RBPMS confers VSMC-specific splicing decisions in HeLa nuclear extract

Overexpression of RBPMS in HEK293 cells induces skipping of *Tpm1* exon 3, dependent on tandem CAC clusters in the flanking intronic regions (Nakagaki-Silva et al., 2019). To further examine the effects of RBPMS on *Tpm1* exon 3 splicing, we employed cell-free *in vitro* splicing assays with two different radiolabeled substrates, TM2-3-4 (Gooding et al., 1994) and TM1-3-4 (Wollerton et al., 2001). The TM2-3-4 substrate comprises *Tpm1* exons 2, 3, 4 and essential flanking intronic regulatory sequences (Fig. S5A). Since *Tpm1* exons 2 and 3 are mutually exclusive, TM2-3-4 provides a simple, binary 5’ splice site choice where exon 4 can be spliced to either exon 2 (2:4) or exon 3 (3:4). In HeLa nuclear extract (NE), TM2-3-4 default splicing generated approximately equal amounts of 2:4 and 3:4 products (Fig. S5B, lane 6). Titration of RBPMS led to a loss of the 3:4 product and a reciprocal increase in the 2:4 product, consistent with repression of exon 3 by RBPMS (Fig. S5B, lanes 5-1).

The TM1-3-4 substrate, comprising *Tpm1* exons 1, 3 and 4, showed even more emphatic changes in splicing outcome upon RBPMS titration (Fig. 2A). The default splicing pattern of TM1-3-4 in HeLa nuclear extract is exon 3 inclusion, with bands corresponding to fully spliced 1:3:4 and partially spliced 1:3-4 and 1-3:4 intermediates (empty circles, Fig. 2A). Only a very small amount of the exon skipping product (1-4) and the corresponding lariat is seen (black filled circles, Fig. 2A). Strikingly, titration of FL RBPMS led to a complete switch from exon 3 inclusion to skipping, indicating that RBPMS is sufficient to confer SMC-specific splicing *in vitro* by interfering with *Tpm1* exon 3 definition (Fig. 2A, lanes 6-2). When tandem CAC clusters both upstream and downstream of exon 3 were mutated, the basal pattern of splicing was unaltered but RBPMS no longer mediated exon 3 skipping (Fig. 2B). Furthermore, the RBPMS ΔC20 mutant failed to induce exon 3 skipping even with intact flanking CAC clusters (Fig. 2C), consistent with observations *in vivo* (Fig 1C). The cell-free AS assay therefore faithfully reflects the specificity of cell culture assays.

**Figure 2.**
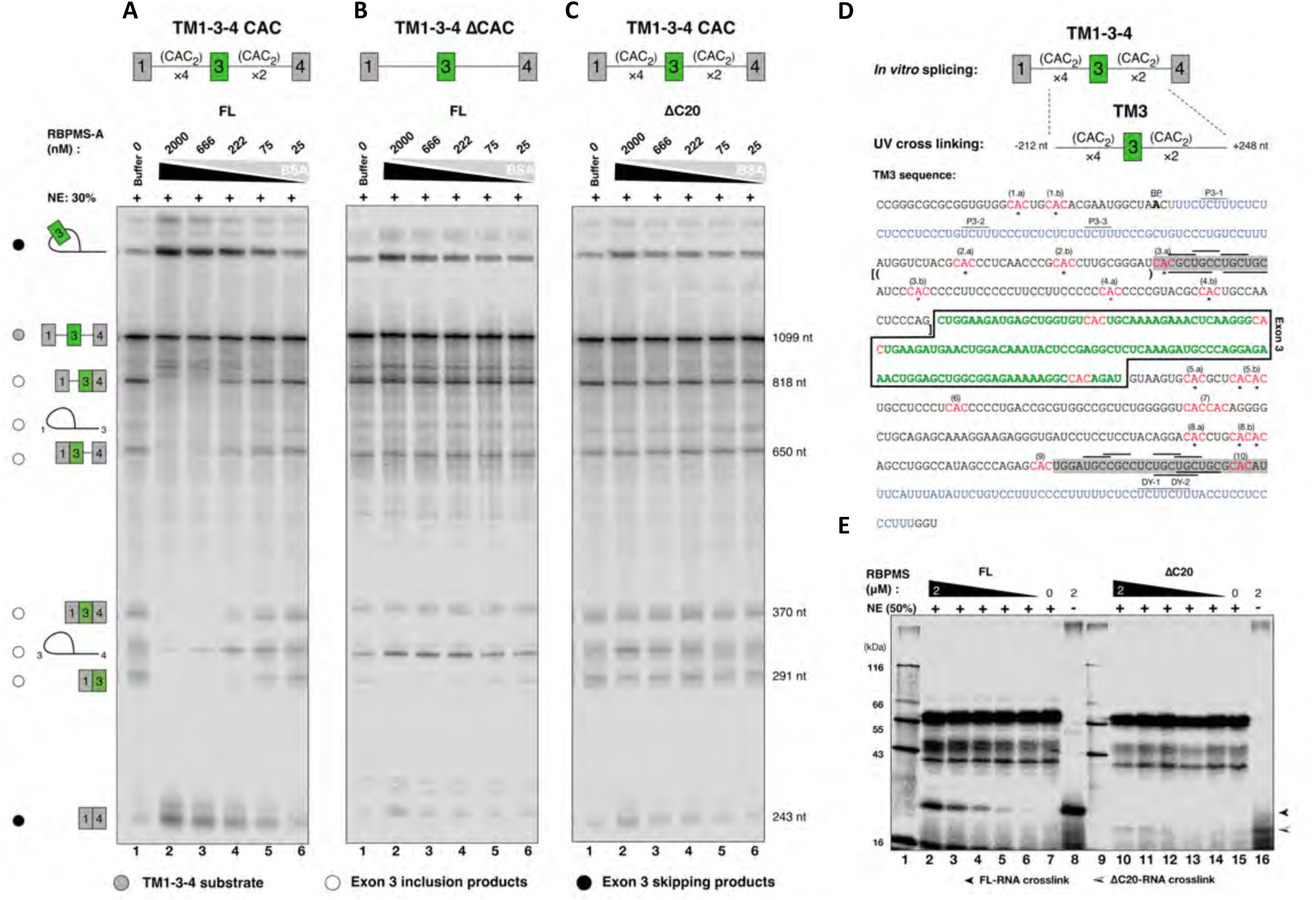
Concentration-dependent RBPMS-A modulation of TM134 *in vitro* splicing reactions. (A) Titration of triply diluted series of FL RBPMS-A into the *in vitro* splicing reaction using TM134 substrate with flanking tandem CAC clusters, (B) using flanking CAC clusters deleted or mutated as specified in Table S3. (C) Titration of ΔC20 into the *in vitro* TM134 splicing reaction. The identities of the linear spliced products are inferred by nucleotide length and depicted by schematic diagrams. The identities of the lariat are determined by matching against the previous *in vitro* splicing experiments (Wollerton *et al* 2001) of the same substrate. The pre-mRNA substrate TM134 is indicated by a grey filled circle. 1-3-4 splicing products are indicated by opened circles, while black filled circles are placed aside of the 1-4 splicing products. One representative of three technical repeats is shown. (D) Top, the TM134-derived TM3 model RNA, containing the full complement of the flanking regulatory sequences. Bottom, the sequence of the TM3 model RNA. Exon 3 sequence is colored in green and boxed. CAC sites are highlighted in red and numbered. C to A mutation or deletion of adenosine in αCAC construct was indicated by black or red asterisks. The exon 3 branch point is indicated by the bolded “A”. Upstream and downstream polypyrimidine tract, colored blue, contains PTBP binding sites that are labelled as P3-X or DY-X. The previously determined flanking regulatory elements, URE and DRE, are shadowed in grey, within which “YGCY” motifs (overlined) and “UGC” motifs (underlined) are able to bind muscleblind-like protein, MBNL1. The sequence of (CAC)_2_ or 3×(CAC)_2_ EMSA substrate bracketed by () or [], respectively. (E) Binding of the triply diluted series of either FL or ΔC20 RBPMS to tandem CAC clusters in Hela nuclear extract. After UV cross-linked to the radiolabelled RNA and RNase digested, the total protein component was resolved on a 15% SDS-PAGE and imaged via phosphor-imaging. The expected mobility of RBPMS-RNA crosslink is indicated by lanes 8 and 16 where no NE was added. One representative of two technical repeats is shown.

### RBPMS C-terminus is essential for RBPMS cooperative RNA binding

To test whether the inability of RBPMS ΔC20 to mediate exon 3 repression was associated with altered RNA binding, we performed UV crosslinking of RBPMS along with HeLa nuclear extract proteins to a radiolabeled RNA substrate TM3, which contains exon 3 and all flanking splice site and regulatory elements (Fig. 2D). While FL RBPMS efficiently cross-linked to TM3 (Fig. 2E, lanes 7-2), very little crosslinking was observed for the ΔC20 mutant (Fig. 2E, lanes 15-10). Hence, the C-terminal 20 amino acids are critical for RBPMS binding to the regulated *Tpm1* pre-RNA in nuclear extract and the RRM domain alone, which mediates both dimerization and RNA binding *in vitro*, is not sufficient for RNA binding in nuclear extract. We further examined the RNA binding properties of FL and ΔC20 RBPMS using electrophoretic mobility shift assay (EMSA). Strikingly, with an RNA substrate containing a single pair of CAC motifs ((CAC)_2_, separated by 10 nt (indicated by curved brackets in Fig 2D), RBPMS ΔC20 bound with an apparent K_d_ of 2.5 μM, whereas FL RBPMS did not bind within the concentration range assayed (Fig. 3A). In contrast, both proteins bound to a longer substrate with three tandem CAC motifs (3×(CAC)_2_ (indicated by square brackets in Fig 2D), with an apparent K_d_ of ∼150 nM, which is 20-fold higher in affinity than ΔC20 binding to the shorter (CAC)_2_ substrate. Importantly, FL RBPMS showed an additional supershifted species with limited gel mobility, which we postulate to be higher-order RBPMS-bound substrates (Fig.3.B, Bound II). Indeed, a derived Hill factor of 1.7 (Fig. 3D) suggests that FL RBPMS bound the 3×(CAC)_2_ substrate in a cooperative manner, whereas ΔC20 binding was non-cooperative. These data indicate that the C-terminal 20 amino acid tail mediates cooperative binding to multivalent RNA substrates, which could be a result of its propensity for driving RBPMS oligomerization (Fig.1). We postulate that in the presence of competing RBPs in the nuclear extract, cooperative multivalent RNA binding (Fig. 2E) combined with interactions with other RBPs (see below) is a requirement for RBPMS to bind target RNAs effectively.

**Figure 3.**
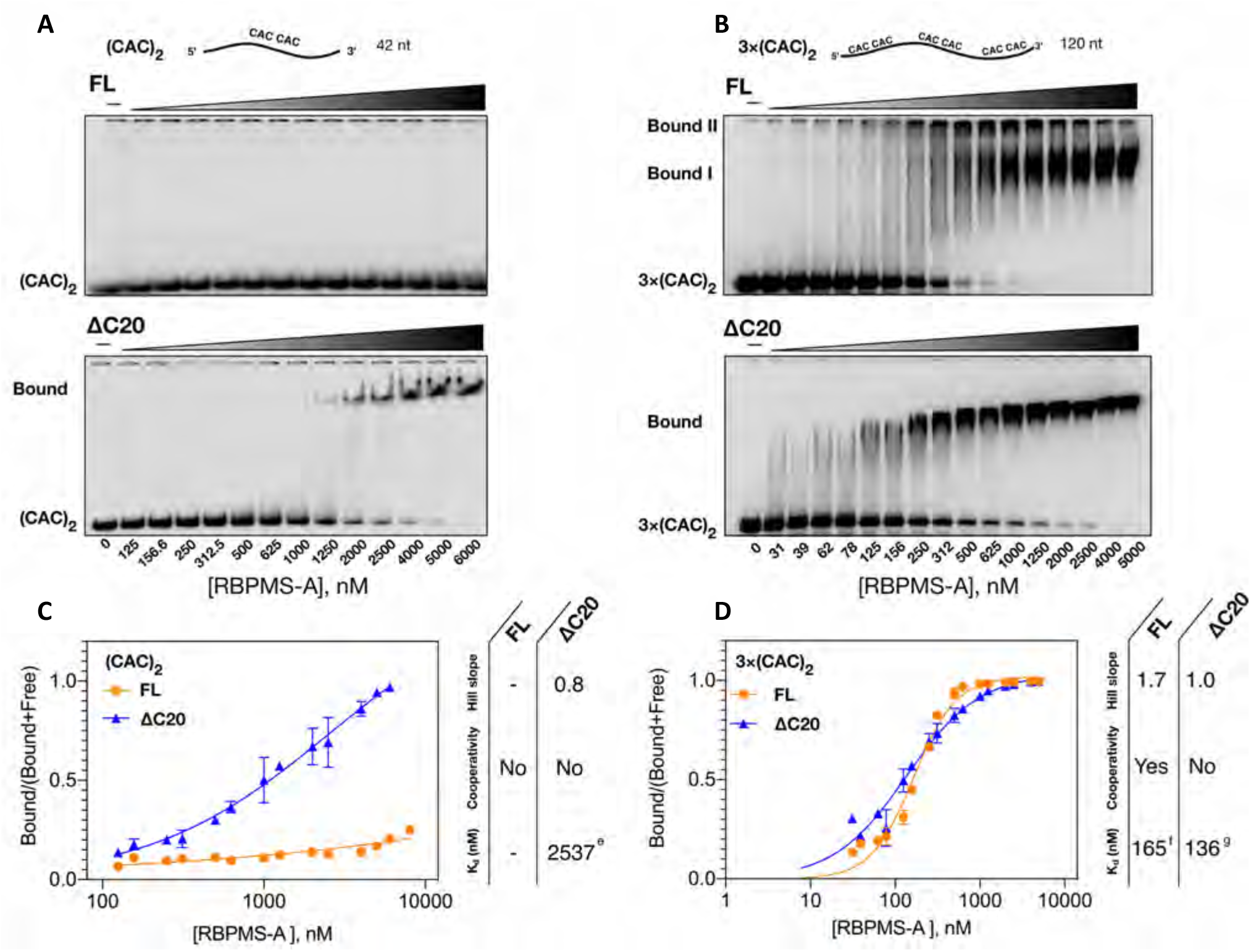
FL and ΔC20 RBPMS-A binding to 10 nM (CAC_2_) substrates in electrophoreCc mobility shiF assay (EMSA). [32p-CTP] incorporated syntheCc RNA of 42 nt, containing single tandem CAC moCf (10 nt spacer Fig 2D), (CAC)_2_, was incubated with 0-6 μM RBPMS recombinant protein as indicated and resolved on a 5% naCve polyacrylamide gel. [^32^P-CTP] incorporated syntheCc RNA of 120 nt, containing three tandem CAC moCfs (10-16 nt spacers Fig 2D), 3×(CAC)_^2^_, was incubated with 0-5 μM RBPMS recombinant protein as indicated and resolved on a 5% naCve polyacrylamide gel. One representaCve of three technical repeats is shown. (C-D) DeterminaCon of dissociaCon constants (K_^d^_) for RBPMS recombinant proteins binding to either (CAC)_2_ or 3×(CAC)_2_ substrates. AFer phosphor-imaging, every lane was quantified as two proporCons, “Bound” and “Free”. The specific binding was determined as Bound/(Bound+Free), which was plo_ed against protein concentraCon. (e) The 95% confidence interval (CI) is between 1257 and 18719 nM. (f) The 95% CI is between 152.4 and 177.7 nM. (g) The 95% CI is between 118.2 and 159.3 nM. The data points and error bars depict the mean of three technical repeats and the standard deviaCon. Data points of protein at 0 nM, were omi_ed due to the log scaled x-axis. The curves were fitted using specific binding with the Hill slope equaCon (see M&M) provided by GraphPad Prism 9 package.

### RBPMS remodels pre-spliceosomal complexes on target RNA substrates

RBPMS may mediate exon 3 repression by direct modulation of pre-spliceosomal assembly pathways. To explore this possibility, we examined complex assembly on two *cis*-splicing incompetent substrates, TM3 and TM23, in HeLa nuclear extract (Fig. 4 and 5). Both TM3 and TM23 contain *Tpm1* exon 3 with the full complement of splice site and regulatory elements, but no flanking constitutive splice sites, thereby allowing us to focus on complex assembly only across the regulated exon 3 region. To ensure that the complexes forming on TM3 and TM23 are functionally relevant, we first tested their activity in *trans*-splicing assays (Wongpalee *et al*, 2016).

**Figure 4.**
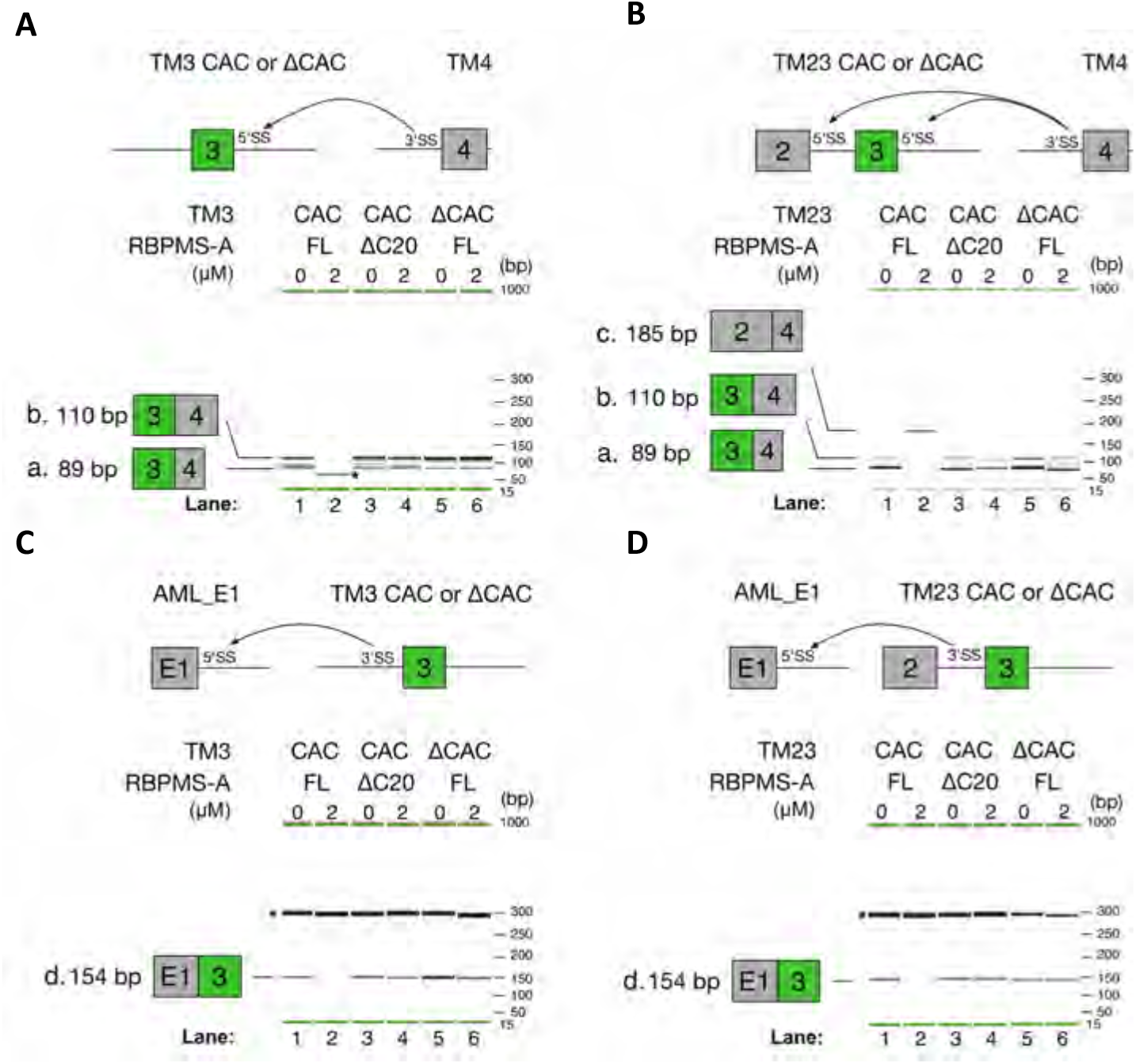
RBPMS regulates 5’ and 3’ SS on TM3 and TM23. The top panels illustrate schematic diagrams showing the *trans*-splicing reactions (black curved arrows) between the TM3 (A) or TM23 (B) and the 3’SS of the TM4 *trans* partner. Sensitivity to FL or ΔC20 RBPMS was tested on either CAC sufficient (CAC) or deficient (αCAC) exon 3 containing substrate. After incubation with NE, total RNA was recovered and reverse transcribed with primer complementary to 3’ end of exon 4. In (A), PCR reactions aimed at amplifying 3-4 *trans* spliced product. QIAxel imaging of PCR product *a*, 89 bp, is the intended 3-4 PCR product. Band *b*, 110 bp, was the result of mispriming of the reverse primer at +20 position with respect to the on-target priming. However, both products report 3-4 *trans* splicing product faithfully. In (B), Three-primers PCR reactions detecting both 2-4 or 3-4 *trans* splicing products were performed. The identical PCR products, *a* and *b*, indicative of 3-4 spliced products were detected, while band *c* appears under FL RBPMS-A modulation (lane 2). Lower panels illustrate the *trans*-splicing reactions (black curved arrows) between the TM3 (C) or TM23 (D) and the 5’SS of the AML exon 1. Sensitivity to *cis*- and *trans*-elements was tested as described above. RT was performed with a primer complementary to the 3’ end of exon 3. In (C-D) The identical PCR product, *d*, indictive of AML_E1-3 *trans* spliced products were detected, but subjected to FL RBPMS-A repression (lane 2). The identifies of band *a*-*d* were confirmed by sequencing. Asterisks were placed next to non-specific PCR amplifications. Extended experiments using concentration series of RBPMS and additional negative controls were shown in Fig S6.

**Figure 5.**
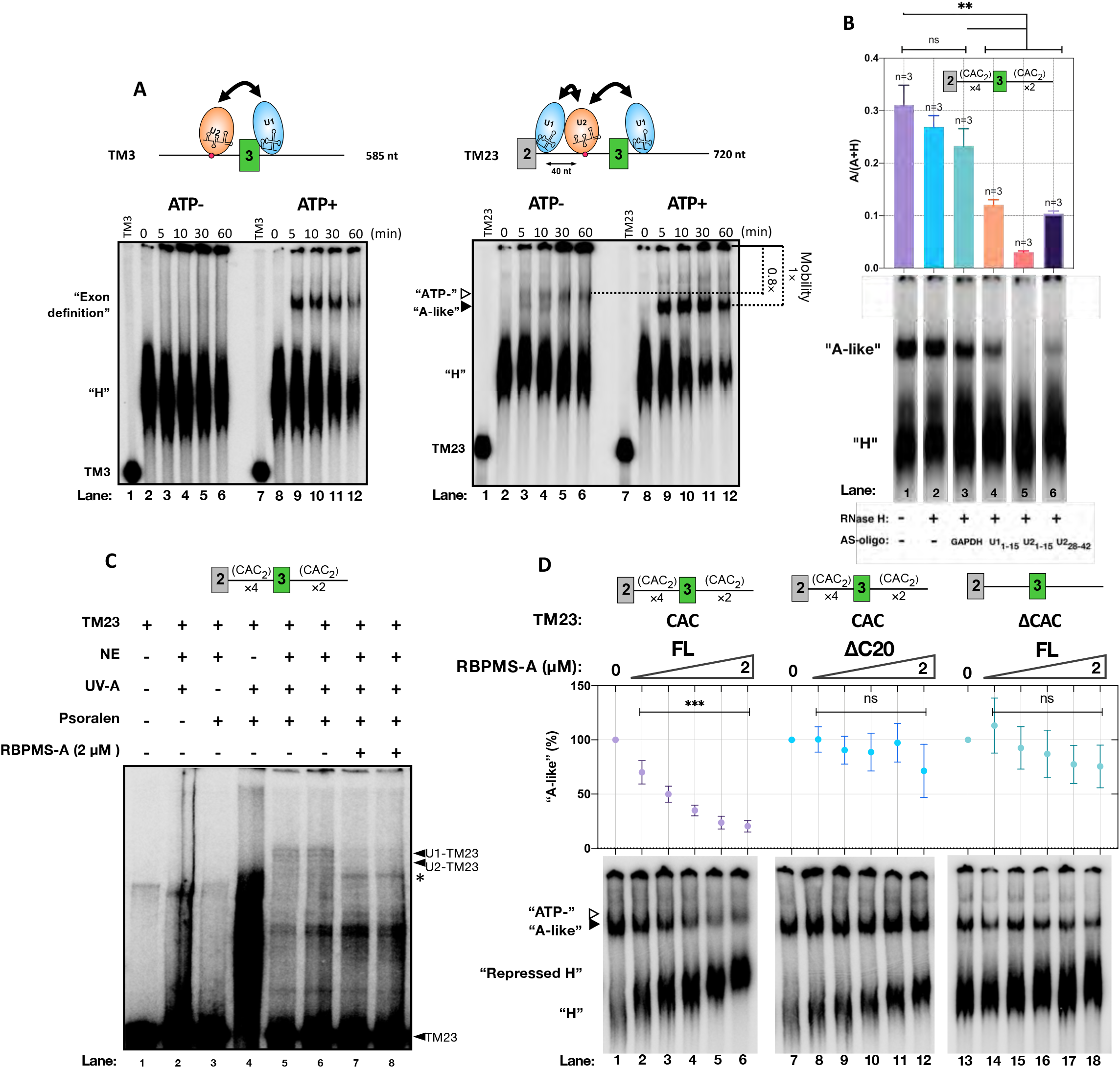
RBPMS modulates pre-spliceosome complexes formed on the miniaturized *Tpm1* exon 3 model RNAs. (A) Two splicing-substrates-derived splicing incompetent model RNAs, TM3 and TM23, were incubated in NE, ±ATP and sampled at the specified time. Branch point sequence denoted by a red dot. Hypothesized binding of snRNPs is illustrated, the potential pairings of snRNPs are indicated by doubled-headed arrows. The mobility position of “ATP-” and “A-like” complexes are indicated by opened and closed triangles, respectively. (B) “A-like” complex formation in sham-treated NE (lane 1), NE pre-treated with RNase H in combination with H_2_O (lane 2) or DNA oligonucleotide complementary to GAPDH mRNA, U1_1-15_, U2_1-15_ or U2_28-42_ snRNA (Lane 3-6). After phosphor-imaging, every lane was quantified as two proportions, “H” and “A-like” complexes, aggregation signals accumulated at the start of the lanes were excluded. The ratio of “A-like” complex conversion was determined as “A-like”/(“A-like”+ “H”), labeled as A/(A+H) on the y-axis. (C) Base-pairing of U1 or U2 snRNA to TM23 in “A-like” complex and in the presence of RBPMS-A. The identities of snRNA-TM23 crosslinks were verified and shown in supplementary Fig S8. Asterisk indicates a possible intra-molecular crosslink that was insensitive to U164-75 or U2104-112 digestion in combination with RNase H, supplementary Fig S10. (D) Incubation of TM23 (CAC) or CAC cluster deficient TM23 (αCAC) in NE in combination with the triply diluted series of FL or ΔC20 RBPMS-A. To determine the “A-like” percentage, the ratio of the “A-like” complex was firstly determined as described in (A). For every titration, RBPMS spiked conditions were normalized against the condition where RBPMS-A was omitted, ((“A-like”_RBPMS_/”A-like”_null_) ×100). For B and D, the data points and error bars depict the mean of three technical repeats and standard deviation. Unpaired, one-tailed, Student’s test was used to assess the differences in mean ratio or percentage of “A-like” conversion across different conditions and are shown as: ns, non-significant; ** p < 0.01 ; ***p < 0.001

To test the 5′ splice site functionality of TM3 and TM23, *Tpm1* exon 4 with an 88 nt 5′ intronic extension (TM4) was used as a 3′ *trans*-splicing partner. TM4 (50 nM) was used at 10-fold molar excess over either TM3 or TM23 (5 nM), in order to overcome splicing inefficiency caused by the physical separation of 5′ and 3′ substrates (Keiper et al., 2019). The 89 nt TM34 spliced product was generated from paired TM3:TM4 and TM23:TM4 reactions in HeLa NE (Band a, Fig. 4A-B, lane 1 and Fig S6). A second band at 110 nt (band b) resulted from mis-priming 21 nt downstream in exon 4, but still reports on TM3:4 splicing. These results confirm that TM3 and TM23 retain 5′ss functionality in *trans* splicing. Moreover, for both *trans* splicing substrates, the addition of FL RBPMS abolished TM34 splicing (Fig. 4A-B, lane 2). With the TM23 substrate, we included a third PCR primer to detect TM24 splicing (185 nt, band c, Fig. 4B). Remarkably, addition of >75 nM RBPMS induced a complete switch from TM34 to TM24 splicing (Fig 4B lane 2, S6B), indicating that RBPMS regulates 5′SS competition between exon 2 and 3. In other words, RBPMS mediated alternative *trans*-splicing *in vitro* (Fig. 4B). This is an important observation that demonstrates the specificity of RBPMS action. The 5′SS of exon 2 is only 41 nt upstream of the exon 3 branch point; the activation of exon 2 splicing demonstrates that RBPMS is not “smothering” the whole RNA to make it splicing incompetent, but is acting in a precise and targeted manner. Again, for both TM3 and TM23, RBPMS splicing regulatory activity was completely dependent on its C-terminal 20 amino acids (lanes 4) and the presence of CAC motifs flanking exon 3 (lanes 6). The identities of bands a, b and c were confirmed by sequencing, and control lanes showed that they only appeared upon incubation with both *trans* spliced partner RNAs and in the presence of ATP (Fig S6, lanes 19-24).

We tried to test the 3′ splice site functionality of TM3 and TM23 using *Tpm1* exon 1 with its downstream intronic segment (TM1) as a 5′ *trans*-splicing partner. We were unable to detect the TM1:3 splice product with either the TM3 or TM23 acceptor (data not shown), even though equivalent splice products were readily detected in *cis* splicing experiments (Fig. 2). We suspected that the large size of the TM1 substrate (350 nt) might result in inefficient *trans-*splicing kinetics. We therefore used the 34 nt Adenovirus Major Late exon 1 with 95 nt of 3′ intronic sequence (AML_E1) (Wongpalee et al., 2016). We detected AML-TM3 splice products at 154 nt, confirming the 3′ SS functionalities of TM3 and TM23 (band d, Fig. 4C-D). Again, RBPMS inhibited the *trans* splicing of AML_E1 to exon 3 for both substrates (Fig. 4C-D, lanes 1,2). This effect was again dependent on an intact RBPMS C-terminal region (lanes 3,4) and the presence of CAC clusters on both sides of the regulated exon (lanes 5,6).

Having established that both TM3 and TM23 are competent for *trans* splicing and are regulated specifically by RBPMS, we proceeded to investigate how RBPMS regulates the assembly of splicing related complexes (Fig. 5). In the absence of RBPMS and with ATP present, both minimal substrates initially formed a heterogeneous (H) complex of fast mobility which developed into a lower mobility complex within 5 minutes (Fig. 5A, ATP+). The lower mobility ATP-dependent complexes were sensitive to targeted partial digestion of U1 and U2 snRNA (Fig. 5A-B, Fig. S7A-B). Psoralen crosslinking further confirmed U1 and U2 snRNA base pairing to TM23 (Fig. 5C, lanes 5-6, Fig. S8). Given the single exon configuration of TM3, we propose that the lower mobility complex on TM3 corresponds to an exon definition A (EDA) complex (Sharma et al., 2008). On the other hand, the advanced complex formed on TM23 could be a combination of an EDA complex across exon 3 and a sterically hindered “A-like” complex between exons 2 and 3 (Smith & Nadal-Ginard, 1989). Notably, on TM23 but not TM3, a lower mobility complex also formed in the absence of ATP. This complex could be distinguished from the ATP dependent “A-like” complex by its slightly lower mobility (∼0.8 fold lower mobility).

Addition of FL RBPMS abolished the formation of ATP-dependent complexes on TM3 and TM23 in a concentration dependent manner (Fig. 5D, Fig. S9A). At the highest concentration (2 μM) of RBPMS approximately 20% of lower mobility complexes remained on TM23 (Fig 5D). However, the residual low mobility complex migrated more slowly than the ATP dependent complex in the absence of RBPMS (∼0.8 fold lower mobility), similar to the ATP-independent low mobility complex (Fig 5A). RBPMS therefore appears to inhibit formation of all ATP dependent complexes. Consistent with this, in the presence of RBPMS, U1 and U2 snRNA base-pairing to TM23 was eliminated (Fig. 5C, lanes 7-8; Fig. S10). Meanwhile, the progressive reduction in H-complex gel mobility is indicative of RBPMS binding (Fig. 5D, left panel). All the effects of RBPMS upon splicing complexes were dependent on the C-terminal 20 amino acids of RBPMS and clusters of tandem CAC sites flanking the regulated exon (Fig. 5D middle and right panels, Fig. S9), mirroring the requirements for RBPMS splicing regulation in *cis* and *trans* splicing assays (Figs. 2 and 4). Taken together, our results establish a strong link between RBPMS splicing regulatory activity and its remodeling of spliceosomal complexes on model transcripts.

### RBPMS remodels the RNA bound proteome composition

Changes in gel mobility of complexes forming on TM3 and TM23 are expected to be caused not only by RBPMS binding and snRNP displacement, but also by the recruitment and displacement of other RBPs. In line with this hypothesis, FL RBPMS was observed to alter the crosslinking of other nuclear extract proteins to the TM3 substrate (Fig. 2E, lanes 2-7). To identify RBPMS binding partners in HeLa nuclear extract, we first generated recombinant RBPMS proteins with an N-terminal Strep-tag II followed by a polyhistidine-tag (StrepII-His_6_-RBPMS) for affinity purification–mass spectrometry (AP-MS) experiments (Fig. S11 and S13, Fig. 6C). StrepII-His_6_-RBPMS has similar oligomerization properties to untagged RBPMS, indicating that the tag has negligible effects on RBPMS biophysical and functional properties (Fig. S4A). After filtering out common contaminants (Fig. S13B), a total of 118 FL RBPMS interactors were significantly enriched above background and 118 FL interactors are significantly depleted from the ΔC20 pulldown (Fig S13C, Supplementary Data File AP_MS_SL). Interactors were then classified using enriched gene ontology (GO) terms on STRING (Szklarczyk et al, 2019). Analysis of both lists of interactors generated near identical top five enriched terms from each category (biological process, molecular function and cellular component) (Fig. 6A and Fig. S13B). 50 RBPMS interactors were selected from the terms: RNA splicing, RNA binding and Ribonucleoprotein complex, grouped and illustrated (Fig. 6B) based on their annotated function as 3′ end processing factors, heterogenous nuclear RNPs (hnRNPs), splicing regulators, other RBPs, interaction network of RBFOX2 (Damianov et al., 2016), RNA helicases, and components of pre-spliceosome complexes such as U4/U6·U5 tri-snRNP and U2 snRNP. Among the splicing regulators was MBNL1, a known regulator of *Tpm1* splicing (Gooding et al., 2013). Remarkably, truncation of the C-terminal 20 amino acids led to a near global loss of the RBPMS interactome (Fig. 6B).

**Figure 6.**
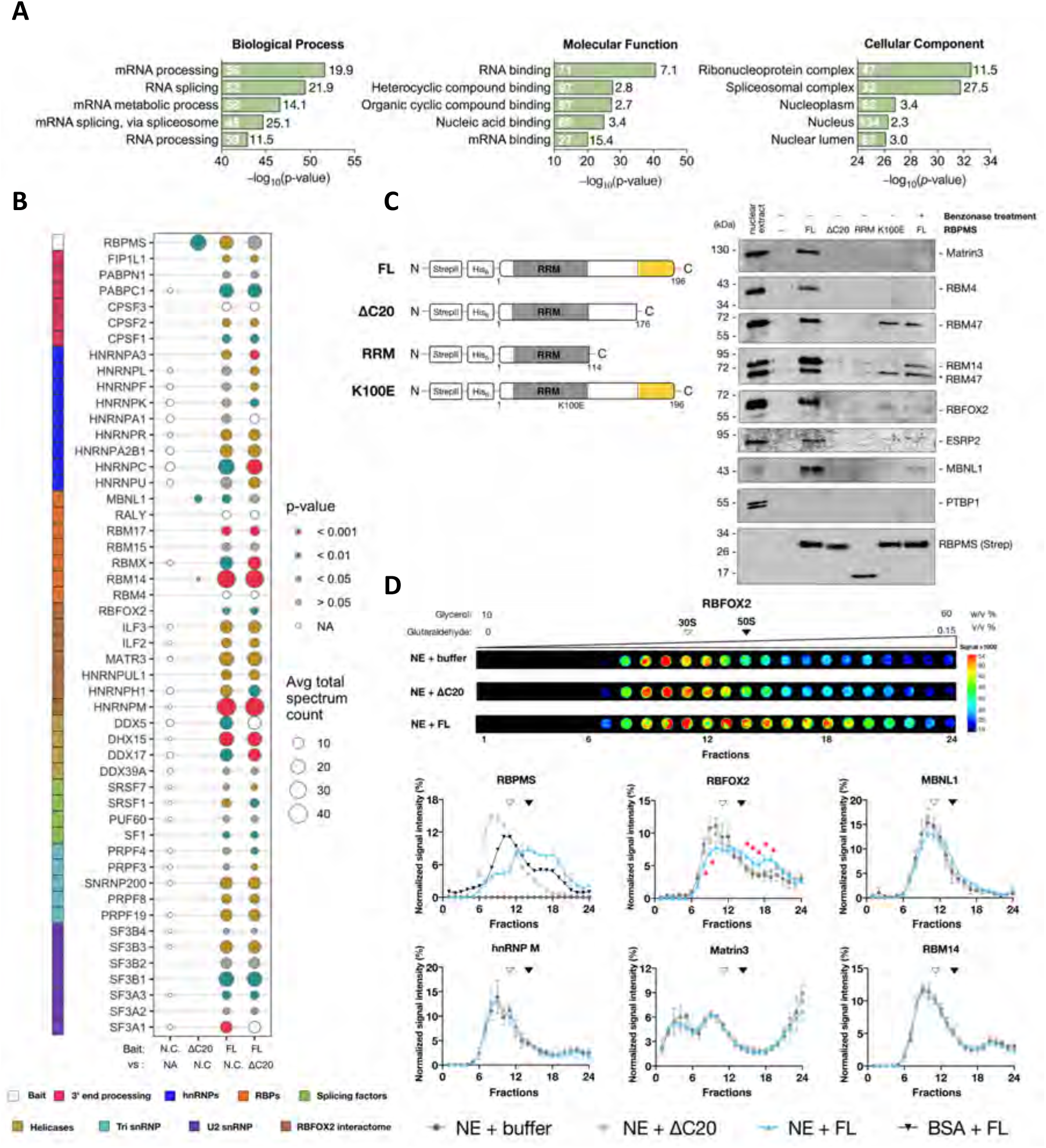
RBPMS-A interactome in Hela NE. (A) GO analysis of the 118 RBPMS-A interactors detectable above negative control. The significance of enrichment for each term was evaluated using the right-sided hypergeometric test with Benjamini-Hochberg *p*-value correction. The top five enriched terms are shown for each GO category (biological process, molecular function, and cellular component). The number within the bar in white indicates the number of proteins annotated with a particular term. The number in front of the bar in black indicates the strength of the enrichment effect (i.e. the ratio between the number of proteins annotated with a term in an enriched network and the number of proteins expected to be annotated with this term in a random network of the same size). (B) A short-listed 50 AP-MS identified interaction partners, the pull-down of which is dependent on the presence of the RBPMS-A C-terminal tail. Avg total spectrum count, the average number of the raw spectral counts across replicates. hnRNP, heterogenous nuclear ribonucleoprotein; RBP, RNA binding protein. Statistical comparisons of the raw peptide count replicates were conducted as specified under the x-axis and by unpaired, one-tailed, Student’s T test. *P*-values are color-coded by the significant levels. Due to the low peptide count and variability within certain replicates, t-tests in these cases were inconclusive, therefore annotated as NA. (C) Left panel, schematic diagram of StrepII and His6 tagged RBPMS variants used. Right panel, Immunoblot assessment of the dependency of RBPMS-A interactome to C-terminal truncations, an RNA binding mutation K100E, and nuclease treatment. Proteins probed are indicated on the right (see M&M for antibodies used). Strep-His_6_-RBPMS-A and its truncations were detected with an antibody to the Strep II tag. (D) Top panel, dot blot analysis of RBFOX2 sedimentation resolved by glycerol gradient loaded with nuclease treated Hela nuclear extract in combination with either buffer, ΔC20 or FL RBPMS. Gradient fractionations performed from top to bottom are ordered from left to right. Scale bar on the right colour coded for the signal intensities. Bottom panel, normalized signal intensity of indicated protein across fractionated glycerol gradient. Every sedimentation profile was determined by performing two-three technical repeats of the gradient separation followed by dot blot and quantification. Intensity of an individual fraction is normalized to the total signal of a given gradient. Sedimentation position of 30S or 50S *E*.*coli* ribosomal subunit is indicated as open or filled triangle. Error bar indicates, SEM, standard error of the mean. Statistical analysis NE+FL vs NE +buffer was performed using unpaired, one-tailed, Student’s T test, indicated as * = p < 0.05.

To validate some of the RBPMS-mediated interactions we performed western blot analysis (Fig. 6C, right), and confirmed interactions between FL RBPMS and MATR3, RBM4, RBM14, RBM47, RBFOX2, ESRP2, and MBNL1. The lack of interaction with PTBP1, another coregulator of *Tpm1* exon 3, serves as a control for the specificity of RBPMS interactions. All the interactions were completely abolished by the ΔC20 deletion, except for RBM14 and RBM47. Using benzonase-treated nuclear extracts or the K100E RNA-binding mutant, most of the interactions were observed to be partially or completely dependent on RNA binding (Fig.6C, lanes 6,7 compared to 3).

We next tested whether RBPMS detectably altered the glycerol gradient sedimentation profiles of a subset of its interactors (Fig. 6D). Upon addition to nuclear extract, RBPMS itself sedimented more rapidly than free RBPMS, indicating that it forms heterogeneous high molecular weight complexes. Consistent with its loss of both homo-oligomerization (Fig 1) and heterotypic protein-protein interactions (Fig 6 B,C) the ΔC20 RBPMS in nuclear extract sedimented in lighter fractions than FL RBPMS (Fig 6D). Strikingly, RBFOX2 shifted into heavier fractions upon addition of FL-RBPMS but not ΔC20 to nuclear extract (Fig 6D), suggesting that the two proteins are components of a common higher molecular weight complex. RBFOX2 is known to be present in the multicomponent benzonase-resistant Large Assembly of Splicing Regulators (LASR) complex (Damianov et al., 2016). The sedimentation profiles of other proteins, including the LASR components MATR3 and hnRNPM, were unaffected by RBPMS suggesting that the RBPMS-RBFOX2 complex is distinct from LASR. MBNL1 appeared to show a slight shift to heavier complexes, but the differences in MBNL1 were not significant between equivalent fractions in the presence or absence of RBPMS.

Having established that the RBPMS interactome includes numerous splicing factors and regulators, we proceeded to examine how RBPMS remodels the composition of splicing-related complexes on splicing substrates tagged with MS2 sites to facilitate affinity purification with MBP-MS2 (Fig. 7A). We initially attempted to use the TM23 substrate, but were unable to achieve purification of specific complexes, in part due to the large size of TM23 (820 nt). We therefore opted for the shorter TM3 substrate and omitted the molecular crowding agent PVA to facilitate comparable recovery of transcripts across different experimental conditions. Urea-PAGE analysis showed a slight increase in RNA recovery in RBPMS-spiked samples, but these differences were within the normalization range of downstream data processing (Fig. S12C). However, assembly of ATP dependent low mobility complexes was negligible under these conditions (Fig S12A). Total proteins from each condition (± ATP, ± RBPMS) were submitted to LC-coupled quantitative label-free MS/MS. Consistent with the complex assembly conditions, very few snRNP proteins were detected even in the absence of RBPMS. However, differential pull-down analysis revealed that a large number of RNA binding proteins were either recruited to (e.g. ESRP2) or displaced from (e.g. SRSF3) the TM3 substrate by RBPMS (Fig. 7B-C). Proteins identified as RBPMS interactors in the AP-MS experiment were also found among both RBPMS-recruited proteins (e.g. RBM4, RBM14, RBFOX2) and RBPMS-displaced proteins (e.g. SRSF7, hnRNPC) (Supplementary Data File RNA_MS_B). The differential recruitment of some proteins appeared to be sensitive to ATP; for example, enrichment of bound ESRP2 and depletion of SRSF7 by RBPMS was only observed in the presence of ATP. We observed no significant changes in the transcript bound levels of PTBP1 and MBNL1, despite the fact that MBNL1 was identified as a direct interactor (Fig. 6) and both proteins are coregulators that bind to sites flanking *Tpm1* exon 3 (Gooding et al., 2013, Gooding et al., 1998).

**Figure 7.**
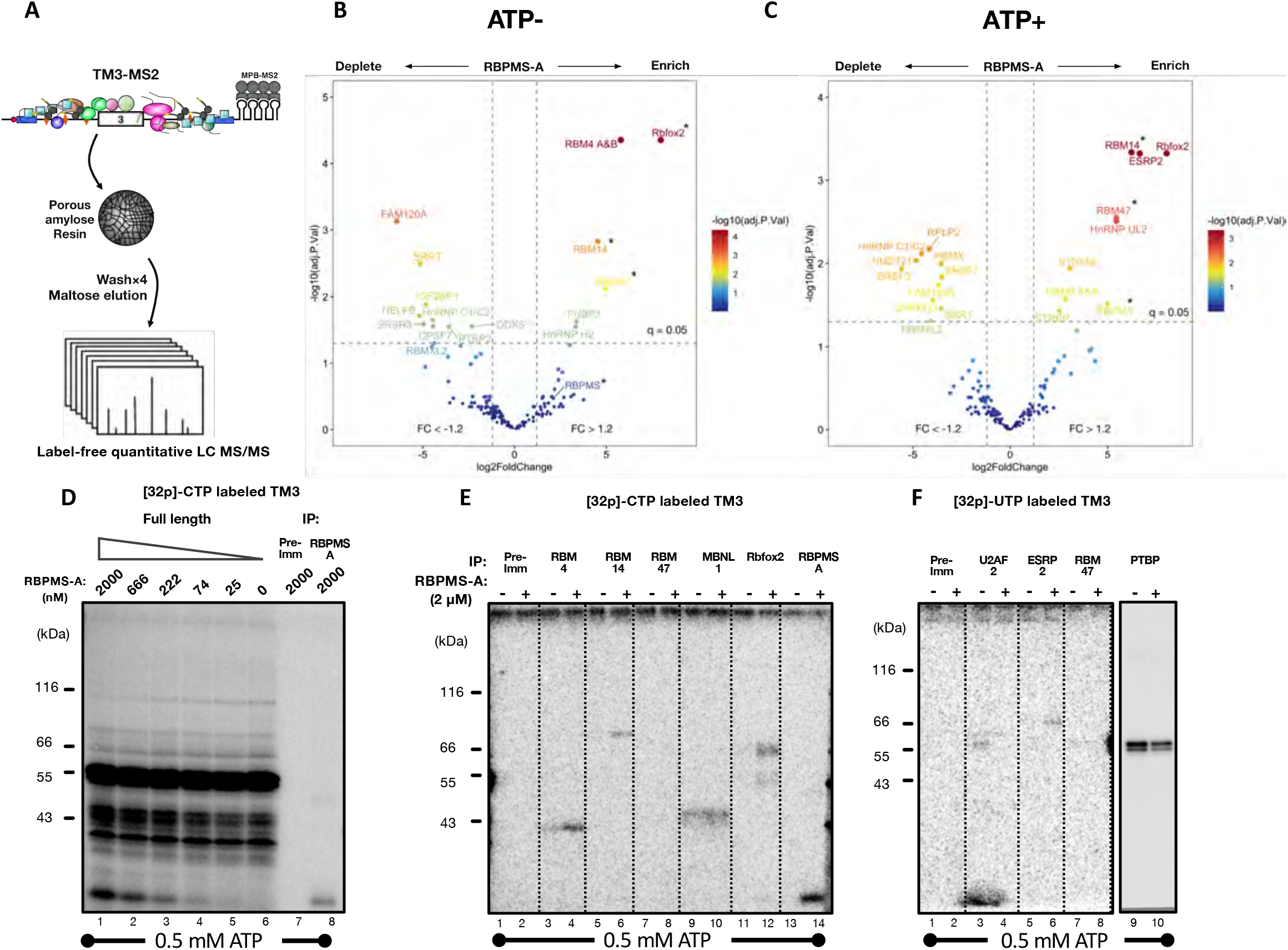
RBPMS-A modulates transcript-bound proteome. (A) Graphic summary of the RNA affinity purification. As described in detail in M&M, MBP-MS2 fusion protein bound TM3-MS2 was incubated in Hela NE under conditions exclusive to “H” complex assembly, ± RBPMS-A (1.5 μM), and ± ATP. The RBPs (coloured shapes) associated with the transcript, were purified via amylose resin, washed, eluted, and subjected to label-free quantitative LC-MS/MS. Volcano plots showing differential pull-down analysis of the proteins discovered from the RNA affinity purification in the absence (B) or presence of ATP (C). Comparisons of ± RBPMS-A conditions gave rise to many proteins either enriched or depleted by RBPMS-A. Asterisk indicates that protein was not detected in the absence of RBPMS-A, the underestimated log2Foldchange was derived using imputed background values. Two vertical lines indicate protein pull-down fold changes estimated by the Bayes method (RBPMS + vs -) >1.2 and <-1.2, respectively. The horizontal line indicates the adjusted *P-value* (False discovery rate, q-value) of 0.05. *P* values were corrected by the Benjamini-Hochberg method. (D) Lane 1-6, Protein binding to radiolabelled TM3 transcript upon titration of FL RBPMS-A was analysed via UV cross-linking and phosphor-imaging. Two immunoprecipitations were performed using samples prepared identically to that shown in Lane 1, with either rabbit pre-immune serum (Lane 7) or rabbit polyclonal antibodies specific to RBPMS (Lane 8).Immunoprecipitations of RNA-protein crosslinks from the digested [32p]-CTP (E) or [32p]-UTP (F) labelled TM3 transcript incubated in NE ± RBPMS-A, in the presence of ATP. Proteins probed are indicated on top using antibody listed in M&M.

Differential RBPMS-sensitive binding of selected RBPs was confirmed by UV cross-linking of [^32^P]-UTP or [^32^P]-CTP labeled TM3 RNA to proteins in HeLa nuclear extract (Fig. 7D-F). RBM4, RBM14, Rbfox2 and ESRP2 crosslinking only occurred in the presence of RBPMS, in agreement with results from the differential pull-down using MS2-tagged TM3. Notably, each of these proteins was seen to interact with RBPMS in an RNA-dependent manner (Fig. 6C). RBM47 crosslinking was not detected with either the [^32^P]-CTP or [^32^P]-UTP labeled transcript, possibly due to poor crosslinking efficiency. In line with the differential pull-down results, crosslinking of MBNL1 and PTBP1 were unchanged by RBPMS (Fig.7E-F, lanes 9-10), Therefore, PTBP1 and MBNL1 binding to TM3 does not require active recruitment, although their transcript-bound activities may still be regulated by RBPMS. Consistent with repression of *Tpm1* exon 3 splicing, crosslinking of the essential splicing factor U2AF2, which recognises the polypyrimidine tract, was reduced by RBPMS (Fig. 7F).

### RBPMS binding partners are functionally involved in the VSMC AS program

The preceding data indicate that RBPMS interacts with and actively recruits a number of RBPs in HeLa NE to TM3 RNA. MBNL1 is an exception as an RBPMS interactor that binds stably to TM3 RNA independent of RBPMS. To test the functional relevance of the identified interactions we performed siRNA mediated knockdowns in PAC1 VSMCs (Rothman et al., 1992) of RBPMS, RBM4 A and B, RBM14, RBM47, RBFOX2, MBNL1 and 2 and ESRP2. All these RBPs are expressed in PAC1 cells, but only RBPMS shows significantly elevated expression in differentiated compared to more proliferative cells (Fig 8A). For most RBPs similar effects on splicing were observed with 2 or 3 independent siRNAs. With ESRP2 significant effects with one siRNA were not replicated with two independent siRNAs, so we did not pursue this protein further. Depletions of the targeted RBPs and mRNAs were verified by RT-qPCR and western blot (Figs 8.B-C, S.14-16). Depletions of RBM 4 (A and B), 14 and 47 had little or no effect on *Tpm1* splicing (Fig S14). In contrast, knockdown of MBNL1/2 and RBFOX2 caused a significant switch from exon 2 to exon 3 inclusion, similar to RBPMS knockdown (Fig 8.D). We also tested whether any of the RBPs regulate other RBPMS repressed (*Actn1*) and activated (*Flnb, Hspg2*) splicing events (Nakagaki-Silva et al., 2019) (Fig 8D). Notably, both MBNL1/2 and RBFOX2 co-regulated all 4 ASEs in the same direction as RBPMS (Fig. 8D), suggesting that they might work widely as RBPMS coregulators of VSMC specific ASEs. Knockdown of the other RBPs had more variable effects (Fig S14). Depletion of RBM4 and RBM14 led to significant changes in some AS events, consistently in the same direction as RBPMS. In contrast, RBM47 acted in concert with RBPMS on *Actn1*, but antagonistically on *Flnb* and *Hspg2*. Overall, our data show that RBPs recruited by RBPMS to *Tpm1* RNA also act as co-regulators of RBPMS-mediated splicing, with MBNL1/2 and RBFOX2 showing consistent effects concordant with RBPMS on all model VSMC ASEs tested.

**Figure 8.**
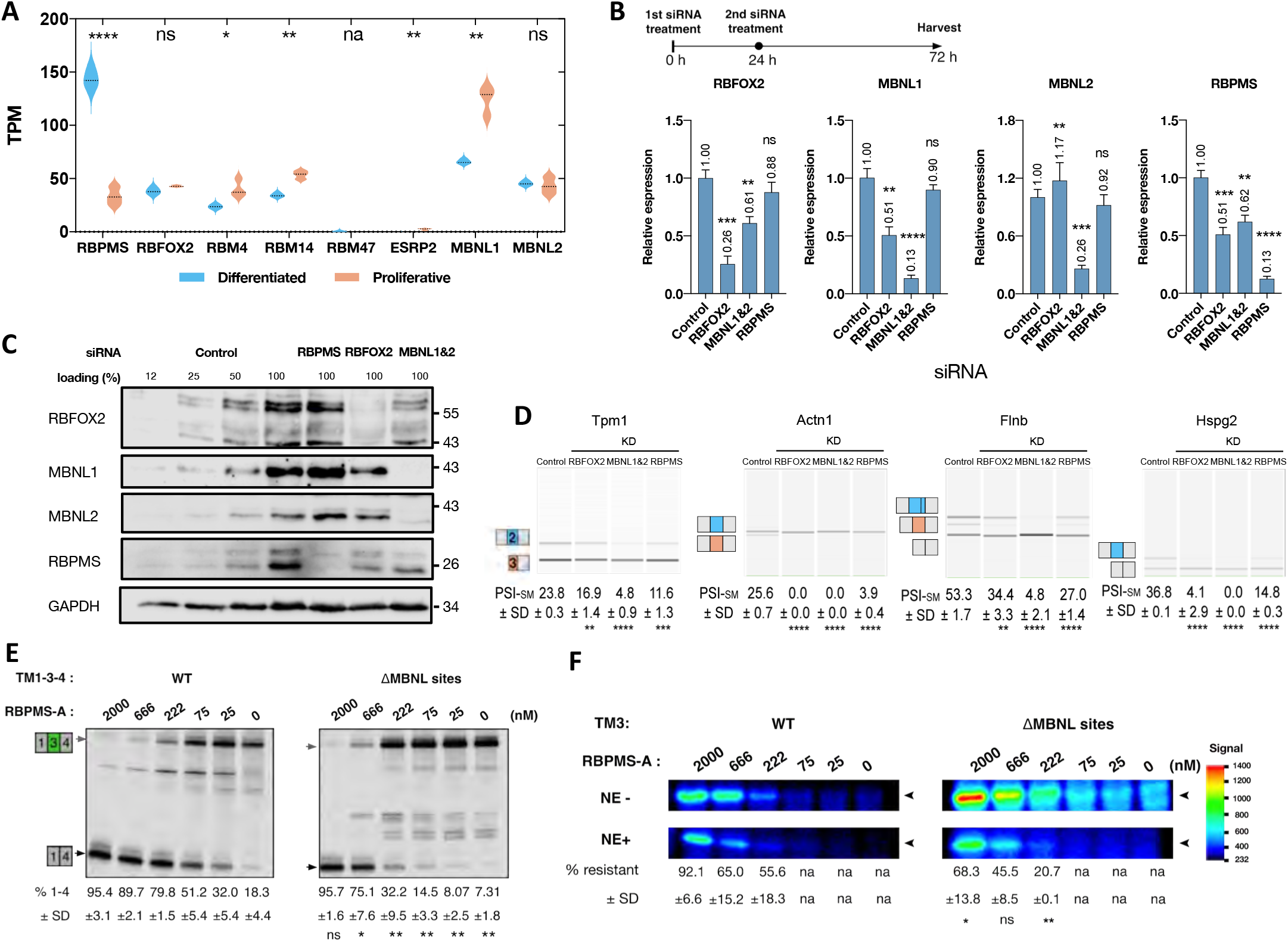
RBPMS-A and its associated proteins modulate VSMC alternative splicing events. (A) mRNA expression of RBPMS and its putative interaction partners in cultured smooth muscle cell PAC-1 in either differentiated, D, or proliferative, P, state. Median of the dataset is depicted as black dashed line. Data retrieved using NCBI Gene Expression Omnibus accession number GSE127799. (B-D) Upon two siRNA treatments, knockdown (KD) of RBPMS, RBFOX2, MBNL1 and MBNL2 was verified via RT-qPCR or Western blots. For qPCR, relative expression levels were normalised to housekeeping gene CANX and Rpl32. Mean and SD are indicated on top of the bar and using error bar. For Western blot, GAPDH detection was used as a loading control. RT-PCR analysis of differentiated and proliferative exon usage upon two siRNA treatments with indicated siRNA. Illustrations of the splicing isoforms are shown on the left and indicate differentiated exon (blue) and proliferative exon (Salmon). PSI, percentage of splice in, values for the SM exons are shown as mean PSI ± SD (n=3). (E) Sensitivity of RBPMS modulated splicing switch on TM1-3-4 to the deletion of flanking MBNL binding sites (URE and DRE elements Fig 2B). (F) Examination of the cooperativity between RBPMS and MBNL1 protein on the binding of [32p]-CTP labelled TM3 RNA using UV crosslinking. %_resistant_, the percentage of binding that sustained NE competition, (Signal_NE+/_Signal_NE-_)100. Statistical analysis was performed using unpaired, two-tailed for A-B and D or one-tailed for E-F, Student’s T test, indicated as ns = p > 0.05, * = p < 0.05, ** = p < 0.01, *** = p< 0.001, and **** = p< 0.0001. For A, RBM47 expression in rat VSMC is insufficient for reliable statistical comparison. For F, signal to noise ratio between 25 -222 nM is too low to conduct confident comparison. siRNA used: RBPMS KD1, MBNL1 THH2, MBNL2, RBFOX2 KD2, see M&M.

MBNL1 regulates *Tpm1* exon 3 splicing by binding to short upstream and downstream YGCY clusters (Gooding et al., 2013). Unlike the other RBPs its stable binding to TM3 RNA was unaffected by RBPMS (Fig 7E). Deletion of both YGCY clusters (ΔMBNL sites, Fig 8E) led to reduced sensitivity to RBPMS regulation of splicing *in vitro*; higher concentrations of RBPMS (4-fold increase in IC_50_) were required to cause skipping of exon 3 in the ΔMBNL site transcript (Fig 8E). Moreover, the reduced activity was also reflected in reduced RBPMS crosslinking to ΔMBNL RNA in nuclear extract, when normalized to crosslinking in the absence of extract (Fig 8F). This suggests that protein-protein interactions with MBNL1 help to recruit RBPMS to TM3 RNA in nuclear extract, thereby explaining the inability of ΔC20 RBPMS to bind to or regulate TM134 RNA (Fig 2).

## Discussion

The activity of recombinant RBPMS *in vitro* allowed detailed analysis of the relationship between its biophysical properties and splicing regulatory activity and insights into how cell-specific splicing regulators interact physically and functionally with more widely expressed RBPs (Fig 9). *Tpm1* exon 3 is efficiently spliced in most cell types, and in HeLa nuclear extract *in vitro*, despite the binding of up to 6 PTBP1 and 3-8 MBNL co-repressors around the exon (Fig 9A). RBPMS exists as a heterogeneous dynamic mixture of dimeric and oligomeric species (Fig 9B, left), with the C-terminal IDR mediating both homomeric oligomerization and heterotypic interactions with other proteins. Oligomeric RBPMS can therefore make multivalent interactions with the multiple (CAC)_2_ motifs flanking *Tpm1* exon 3 as well as contacting other RBPs, which might further stabilize RNA binding. Notably, MBNL1 binds independently to YGCY motifs and by a direct protein-protein interaction helps to recruit RBPMS to the RNA. RBPMS in turn recruits further co-regulators, including RBFOX2, that do not have specific binding sites. As a result a stable repressed complex forms that prevents splicing complex assembly including binding of U1 and U2 snRNPs (Fig 9B right). With deletion of the C-terminal 20 aa of the IDR, RBPMS exists only as a dimer, is unable to interact with other RBPs and consequently is inactive as a splicing regulator, being unable to promote regulatory complex assembly (Fig 9C). We propose that the stable repressed complex, which encompasses a 500 nt region surrounding exon 3 (Fig 2D) resembles the “binding region condensates” described by (Hallegger et al., 2021). Single molecule analyses showed that the TM3 RNA binds ∼5-6 PTBP1 and 3-8 MBNL1 molecules (Cherny et al., 2010, Gooding et al., 2013). Equivalent analyses of RBPMS binding has not been carried out, but the size of the RNA-free RBPMS oligomers (Fig 1) suggests that the repressed complex likely contains 5-10 RBPMS dimers, meaning that the size of the RBPMS:MBNL1:PTBP1 complex would be ∼1MDa or larger, without taking into account RBFOX2 and other RBPs that do not have specific binding sites around exon 3. Despite this size, the repressive mode of action must be very precisely targeted because the 5’ splice site of exon 2, only 41 nt upstream of the exon 3 branch point, is activated by the repressive mechanism operating on exon 3, even on a *trans* splicing substrate (Fig 4, Fig S5). It seems plausible that the “zone of repression” is limited by the two PTBP1 binding tracts, which flank the upstream and downstream MBNL and RBPMS binding sites (Fig 2D).

**Figure 9.**
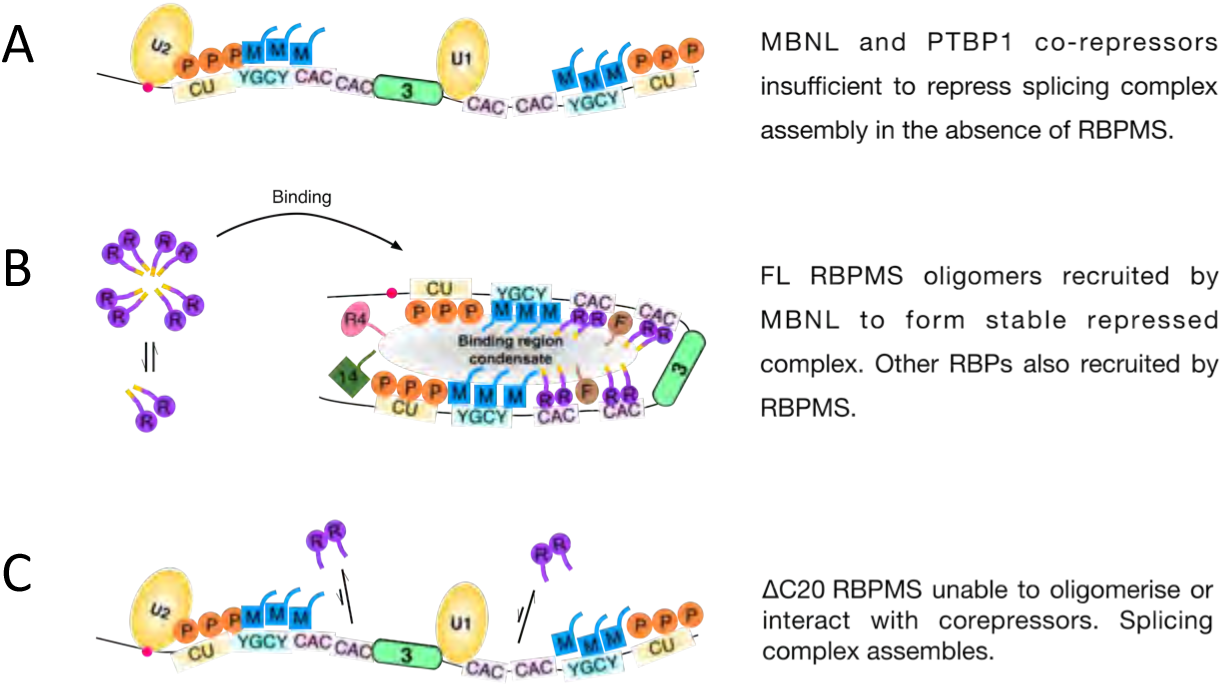
*Graphic summary.* RBP binding motifs, Branch point (Red dot) and *Tpm1* exon 3 were indicated in the strand of RNA (Black line). snRNPs were depicted by yellow ovals. Colored shapes were used to indicate various BRPs, PTBP1 (P), MBNL (M), RBPMS (R), RBFOX2 (F), RBM4 (R4) and RBM14 (14).

Recombinant RBPMS primarily exists as a heterogeneous dynamic mixture of dimeric and oligomeric species, characteristic of phase-separating proteins below their critical saturation concentration (c_sat_) (Kar et al., 2022, Mittag & Pappu, 2022) (Fig. 1D), although it can undergo phase separation forming liquid-like droplets *in vitro* (Figs. 1J, S4B). While the reported association of RBPMS and RBPMS2 with cytoplasmic granules may involve condensate behavior (Farazi et al., 2014, Hörnberg et al., 2013, Kaufman et al., 2018), we envisage that as a splicing regulator RBPMS is in the form of dynamic coregulator-containing hetero-oligomers smaller than the mesoscale assemblies that form visible cellular condensates. The disordered 20 amino acid C-terminal tail, enriched in aromatic and basic residues, is essential for RBPMS oligomerization (Fig 1), alternative splicing outcomes (Figs 1, 2, 4), cooperative binding to multivalent RNA (Figs 2, 3), splicing complex regulation (Fig 5) and most protein-protein interactions (Fig 6). These effects of the ΔC20 deletion are consistent with, and provide a physical basis for, previous reports that C-terminal truncation of RBPMS or RBPMS2 impaired localization to cytoplasmic granules (Hörnberg et al., 2013), participation in RNP complexes (Aguero et al., 2016), co-immunoprecipitation with its full-length counterpart, and mRNA binding (Gerber et al., 2002).

The relative importance of the homomeric and heteromeric interactions mediated by the IDR remains an open question. Indeed, it is plausible that homo- and heterotypic interactions share a common physical basis – for example π-π or cation-π interactions mediated by aromatic and/or basic residues (Wang et al., 2018) — so mutants to distinguish their roles might be elusive. Nevertheless, given the multiple CAC motifs around *Tpm1* exon 3 the ability of RBPMS to oligomerize and cooperatively bind RNA appears to be essential for its function. Likewise, the ability to interact with MBNL1 appears important for recruiting RBPMS to TM3 RNA in the face of competitive binding in nuclear extract (Fig 8). Modulation of RBPMS activity *via* de-oligomerization also appears to be a physiological control mechanism. RBPMS is phosphorylated at Thr 113/118 immediately downstream of the RRM, and phosphomimetic mutants have reduced activity and RNA binding, which is related, in part, to reduced oligomerization, as well as direct occlusion of the RNA binding surface of the RRM in a phosphomimetic mutant (Barnhart et al., 2022).

While many new RBPMS interactors were identified by affinity pulldown (Fig 6), ten proteins in our dataset are known RBPMS interactors including RBFOX2, MBNL1 and RBM14 (Huttlin et al., 2021, Lang et al., 2021, Rual et al., 2005, Yang et al., 2018). That most interactions are direct but enhanced by the presence of RNA (Fig.6.C) is consistent with combinatorial models of splicing regulation, in which the low affinity of binary protein-protein interactions is tuned so as to enable specific cooperative assembly only upon regulated substrates with the correct combination of binding sites (Smith & Valcárcel, 2000). Given the heterogeneity of RNAs in nuclear extract the captured RBPMS interactome is likely to contain RBPMS co-regulators involved in both splicing activation and repression as well as other activities such as 3′ end processing (Fig.6.A). Indeed, the identification of numerous U2 snRNP components suggests possible mechanisms for RBPMS splicing activation by U2 snRNP recruitment. However, it is less clear how this interaction could be involved in the observed displacement of U2 snRNP from *Tpm1* transcripts (Fig 5, Figs S9, S9), which is more likely explained by the displacement of U2AF2 by RBPMS (Fig 7F). In examining how RBPMS remodels the Tpm1 RNA bound proteome, we would ideally have used similar conditions to those used to identify ATP-dependent complexes on native gels (Fig 5D). However, we encountered insurmountable technical problems in trying to purify complexes assembled in the presence of PVA and with the longer (820 nt) TM23 substrate. In the future it would be to useful to exploit single molecule methods to assess how RBPMS affects binding of individual snRNPs to the TM RNAs and also how many RBPMS subunits are associated with the repressed complex (Chen et al., 2017, Cherny et al., 2010). Nevertheless, by analysing complexes formed on TM3 RNA in the absence of PVA, we identified potential splicing coregulators of RBPMS (Fig 7) many of which were also pulled down directly by RBPMS from HeLa NE. In contrast, several known splicing regulators detected in the AP-MS experiment were either depleted from TM3 by RBPMS (e.g. SRSF7 and hnRNP C) or not significantly enriched (e.g. SRSF1). Some of these differences could be attributed to the presence or absence of specific *cis* elements in the TM3 substrate which may be a requirement for recruitment of some interactors, such as RBM4 whose interaction was completely RNA dependent (Fig 6C). RBM4 was previously reported to promote Tpm1 exon 3 inclusion, antagonizing the activity of PTBP1 (Lin & Tarn, 2005), but we saw no effects upon RBM4 knockdown (Fig. S14).

Among RBPMS interactors, we identified many components of the 55S benzonase resistant LASR splicing regulatory complex (Damianov et al., 2016) (Fig. 6C). LASR confers the RNA binding specificities of its other constituent proteins upon RBFOX, effectively expanding the motif recognition preference of RBFOX, consistent with the lack of identifiable RBFOX motifs associated with many RBFOX CLIP tags (Begg et al., 2020). Nevertheless, the higher-order benzonase-resistant RBPMS interaction with RBFOX2 (Fig 6D) did not involve other LASR complex components (e.g. MATR3 or hnRNP M) so it may be a distinct complex. RBFOX2 has been shown to direct distinct splicing outcomes of opposing biological activities by partnering with different splicing regulators (Zhou et al., 2021). RBPMS may have such a determining influence, redirecting RBFOX2 from promoting mesenchymal (Venables et al., 2013) to differentiated VSMC splicing programs. One interesting example is the *Flnb* H1 exon, encoding a hinge region in filamin-B, which is activated in PAC1 cells by RBPMS, RBFOX2 and MBNL1 (Figs 8, S14-16) (Nakagaki-Silva et al., 2019). However, in human breast cancer cells RBFOX1 promoted skipping of the same exon, as part of epithelial-mesenchymal transition (Li et al., 2018).

Several lines of evidence converge upon MBNL1 and RBFOX2 act as general co-regulators with RBPMS. Both proteins were found as a direct RNA-stimulated interactors with RBPMS (Fig 6), RBFOX2 was recruited to the TM3 RNA by RBPMS (Fig 7), while MBNL1 bound TM3 independently via its own binding sites and helped to recruit RBPMS in nuclear extract (Fig 8). Knockdown of both proteins affected four tested ASEs in the same direction as RBPMS (Fig 8). Other lines of evidence also suggest widespread functional cooperation of RBPMS and RBFOX proteins including: enrichment around RBPMS-regulated exons not only of RBPMS dual CAC binding motifs but also of RBFOX binding GCAUG motifs (Jacob et al., 2022); the identification of a GCAUG containing motif as the top intronic binding site for RBPMS in ES cells (Bartsch et al., 2021); and the association of both RBFOX and RBPMS binding motifs with ERG (E26 related gene protein)-repressed exons in HeLa cells (Saulnier et al., 2021). Furthermore, the SMC transcription factor MYOCD was recently found to indirectly drive a set of SMC AS changes via changes in the expression levels of RBPMS, RBFOX2 and MBNL1 (Liu et al., 2022), and both *Rbpms* and *Mbnl1* were found by scRNA-Seq to be part of a contractile VSMC gene signature (Dobnikar et al., 2018). Indeed, it has been suggested that gastrointestinal dysfunction in myotonic dystrophy is associated with dysregulation of an MBNL1 regulated splicing program in visceral smooth muscle cells (Peterson & Cooper, 2022). Our data suggest that this dysregulated program is likely driven by MBNL1-RBPMS coregulation. The finding that recombinant RBPMS is sufficient *in vitro* to switch *Tpm1* exon 3 alternative splicing to the fully differentiated VSMC state (Figure 2), is consistent with its proposed role as a master regulator of the AS splicing program in differentiated VSMCs (Nakagaki-Silva et al., 2019). *Rbpms* heterozygous knockout mice have no phenotype while homozygous knockouts are inviable (Gan et al., 2022) but have phenotypes associated with dysfunction of both VSMCs and cardiomyocytes. Confirmation of the physiological roles of RBPMS in fully differentiated VSMCs *in vivo* will therefore require conditional knockout models.

In conclusion, this study builds on previous work to suggest that the dynamic regulation of splice site choice is dependent on the existence of both constitutive and tissue-specific AS regulatory networks. The intricate connections and functional redundancy of the two networks may reflect the requirement of VSMC to conduct phenotypic switching rapidly in response to environmental cues.

## Materials and methods

### Cloning

The cloning of rat RBPMS-A cDNA, NCBI accession code: XM_006253240.2, into the pEGFP-C1 vector was described previously (Nakagaki-Silva et al., 2019). To produce the RBPMS-A protein with a N-terminal removable His6 tag, PCR products of pET15b vector and TEV-RBPMS-A generated by primers (Table S.1) were treated with T4 DNA polymerase and joined by ligation independent cloning. Based on the resulting pET15b-TEV-RBPMS-A plasmid, PCR product generated by primers (Table S.1) was used to replace the full length RBPMS open reading frame (ORF) flanked by SalI and XhoI restriction sites, producing pET15b-TEV-RBPMS-A-ΔC20 plasmid. To produce RBPMS constructs for affinity purification, the segment of pET15b-TEV-RBPMS-A flanked by XbaI and SalI sites was replaced with DNA oligonucleotide (Table S.2) encoding ribosome binding, Strep-II tag, and His6 tag sequences. Using the resulting pET15b-StrepII-His6-RBPMS-A, restriction enzyme cloning was conducted replacing the full-length RBPMA sequence with either ΔC20 or RRM (2-114 aa) sequences. Also as tabulated in Table S.2, Gibson assembly (Gibson et al., 2009) was used to produce pET15b-StrepII-His6-RBPMS-A-K100E.

### Expression and purification of recombinant protein

Expression vectors were transformed into *Escherichia coli* BL21 (DE3) competent cells. An overnight primary culture was prepared by inoculating a single colony in 10 ml lysogeny broth (LB), 100 mg/ml ampicillin, at 37 °C with shaking. The primary culture was subsequently scaled up by using a 1:50 dilution with LB (100 mg/ml ampicillin), protein expression was induced with 0.2 mM IPTG at O.D.600nm of 0.8 for 2 hr at 37 °C. The post-induction culture was harvested by centrifugation at 7,000*g* for 10 min, the pellet was resuspended in HisA buffer (50 mM Tris, 500 mM KCl, 40 mM Imidazole, 10% v/v glycerol, 0.5 mM DTT pH 8.5). The resuspended cell pellet was lysed using a French press. Nucleic acid precipitation was carried on ice in HisA supplemented with 1M LiCl and cOmplete protease inhibitor (Roche) for 10 min. Lysate was clarified by centrifugation (40,000*g* for 30 mins), the supernatant was filtered with 0.45 μm filter, loaded on a 1ml Histrap HP column (Cytiva) with an AKTA purifier (Cytiva) and eluted with a gradient of increasing imidazole concentration. The identified peak fractions were buffer exchanged into QA buffer (20 mM CAPS, 50 mM KCl, pH10), protein concentration is determined via UV 280nm absorbance. To 1mg of recombinant protein, 25 μg of TEV protease was added and incubated at 4°C for 16hr. The tag removed protein was purified further via a mono Q 5/50 GL column (Cytiva). The NaCl gradient eluted RBPMS-A was pooled and polished with a Superdex 200 16/600 column (Cytiva).

### Analytic ultracentrifigation

Sedimentation velocity (SV) experiments were conducted using an Optima XL-1 analytical ultracentrifuge (Beckman Coulter). Samples were loaded into standard double-sector cells, 12 mm centerpiece thickness, and analyzed at a speed of 40000 rpm with a four-hole An60 Ti rotor, at 20 oC for 15 hours and 300 scans of interference optics were recorded in 90 sec interval. All AUC experiments were performed in buffer containing 20 mM HEPES, 1 mM TCEP, at pH 7.9, but varied in KCl concentrations. Under 500 mM KCl condition, equal molar concentration of full-length and ΔC20 RBPMS were studied at 0.46 and 0.57 mg/ml, separately. At 60 mM KCl, a concentration series of full-length RBPMS was analyzed from 0.7, 0.48, 0.24 to 0.11 mg/ml. Analysis of ΔC20 was conducted at either 0.5 or 0.1 mg/ml. SV data analysis was performed using SEDFIT (v14.1) program, assuming sedimentations of all species fit into a continuous c(s) model. The partial specific volume of the protein (FL _ = 0.73 mL/g, ΔC20 _ = 0.74 mL/g) and the viscosity and density of the buffer (ρ = 1.016 × 10-2; ρ = 1.026) was calculated using the program SEDNTERP (Laue et al., 1992). Best c(s) fits were determined using over 60 scans, by fixing the meniscus, partial specific volume, solvent density, but floating the frictional ratio f/f_0_, until the overall root mean square deviation (RMSD) fall between 0.005 and 0.02. f/f0 between 1 and 1.15 (Fig.S2A-D) was determined to be the compromised value that was used to describe both RBPMS dimer and oligomers in a single c(s) plot. f/f0 values above 1.4 was reached for fitting the FL RBPMS-A at 5 μM (Fig.S2E) and all ΔC20 sedimentation profiles (Fig.S2F-H).

### Cryo-electron microscopy

The sample was spotted on a Quantifoil 1.2/1.3 300 mesh Cu (10) grid (Agar Scientific), blotted, plunge frozen using an Vitrobots (ThermoFisher Scientific). Image acquisition was carried out at the normal magnification of 92,000× using a Falcon 3 counting detector in a Talos Arctica transmission electron microscope (ThermoFisher Scientific). The detailed data collection parameters can be found in Supplementary file EM data collection.

### Fluorophore labelling

Purified His_6_-TEV-RBPMS-A was exchanged to 500 mM KCl AUC buffer using a Zeba spin desalting column (7K MWCO, ThermoFisher Scientific). Alexa Fluor 647 C2 maleimide (ThermoFisher Scientific) was added to 10× molar excess and the labeling reaction was incubated overnight at 4°C in the dark. The reaction was quenched by addition of excess β-mercaptoethanol, and buffer exchanged again to QA buffer. Using an Amicon Ultra centrifugal filter (10K MWCO; ThermoFisher Scientific), labeled protein was repetitively concentrated until a 100,000× dilution is achieved. The amount of free fluorophore in the mixture was estimated by SDS-PAGE (Fig. S4A).

### Phase separation assays

Purified RBPMS-A was exchanged to image buffer (20 mM CAPS, KCl, 1 mM TCEP and pH 10) using a Zeba spin desalting column (7K MWCO, ThermoFisher Scientific). The protein was diluted such that the final concentration was as indicated. For fluorescence microscopy, the mixture included 0.5 μM labeled His_6_-TEV-RBPMS-A in QA. Phase separation was induced by addition of 100 mM HEPES pH 7.9 to final volume of 10 μL. Additionally, PVA was added in experiments using tag-free RBPMS. The mixture was incubated at room temperature (RT) for one hour in the dark. For microscopy, 5 μL of the mixture was spotted onto a glass slide, covered and sealed. Images were acquired using an Nikon ECLIPSE Ti microscope equipped with a 60× oil-immersion differential interference contrast objective. All images were acquired within five hours of the time at which phase separation was induced.

### Band shift

Recombinant proteins were buffer exchanged to Buffer BS (20 mM HEPES, 100 mM KCl and 0.5 mM DTT, pH 7.9). The 3×(CAC)_2_ and (CAC)_2_ RNAs were transcribed using T7 RNA polymerase (Thermofisher Scientific). A 10 μl binding reaction contains 10 nM RNA, 25 mM HEPES, 100 mM KCL, 2 mM MgCl_2_, 0.625 mM DTT, 0.1 mg/ml BSA, 5% glycerol, and increasing concentrations of recombinant RBPMS, at pH 7.9. After 1 hour incubation at 30 oC, 1 μl heparin was added to a final concentration of 5 mg/ml and a 10 min additional incubation was performed at 30°C. Before gel loading, the binding reactions were chilled on ice and 2 μl 50% (v/v) glycerol was added. Bound and free RNA was separated on a native PAGE gel, 5%, 40:1, acrylamide:bisacrylamide ratio, using TBE running buffer at RT. Gels were then dried and visualized by autoradiography on a Typhoon FLA 9000 (Cytiva). Binding curves were fitted with specific binding with Hill slope analysis, Y=Bmax*X^h^(Kd^h^+X^h^), using Prism 9 program.

### Cell culture and nuclear extract (NE) preparation

Hela S3 cells were cultured in suspension with a 5 Liter T-flask in SMEM (Thermofisher Scientific) supplemented with 10% fetal calf serum (SigmaAldrich), with constant agitation at 80 rev/min at 37 oC. 6-8 liters of Hela S3 culture in log phase of the growth, at a cell density of 5×10^5^ cells/ml, were harvested by centrifugation in a Megafuge (Heraeus) at 2000 rpm, at 4°C for 10 min. The cell pellets were immediately washed twice with ice-cold PBS, in total, 40 times the cell pellet volume. The downstream extract preparation was carried out strictly according to the S10 protocol detailed in (Hartmuth et al., 2012).

### In vitro transcription

Depending on the experiment, either [α-32P] CTP or UTP (Perkin Elmer) labeled RNA transcript (specified in the figure legends) were transcribed from linearized pGEM vectors (Table S.7) with T7 polymerase. To make RNA for *in vitro* splicing, complex assembly, and UV cross-linking assays, GTP to m7G(5’)ppp(5’)G dinucleotide cap analog ratio was kept at 1:8 to ensure high capping efficiency. For EMSA assays, the addition of cap analog was omitted from the in vitro transcription mixture. The reaction mixes were tableted in S.11.

### In vitro splicing

*In vitro* splicing was carried out as in (Gooding et al., 1994). Standard *in vitro* splicing reactions were assembled in 10 μl with 20 fmole [^32^P] labelled RNA transcript, 2.2 mM MgCl_2_, 0.5 mM ATP, 20 mM creatine phosphate (Roche), 16U RiboLock RNasin (Thermofisher Scientific), 12 mM HEPES (pH 7.9), 12 % (v/v) glycerol, 60 mM KCl, 0.12 mM EDTA, 0.3 mM DTT, 2.6% polyvinyl alcohol (PVA) and 30 % (v/v) nuclear extract. RBPMS was added at the indicated concentrations in buffer BS. When the effect of RBPMS concentration is studied, in total 500 ng of protein was added for every reaction, a combination of both recombinant RBPMS and BSA (NEB). Splicing reactions were incubated for the 3 hr. After the reaction, reactions were stopped by performing protease K (PK, ThermoFisher Scientific) digestion. RNA components were phenol extracted, ethanol precipitated, and resolved on 4% denaturing Urea-PAGE acrylamide gel. Splicing products were detected by autoradiography with a Phospho-imaging screen and imaged with a Typhoon FLA9000 (Cytiva) imager.

### Complementary DNA oligo directed RNase H breakdown of U1 and U2 snRNP

The DNA oligonucleotides (Table S.4) complementary to 5’ end (Nucleotides 1-15) of snRNA, to the branch point recognition sequence (Nucleotides 18-42) of U2 snRNA and to GAPDH mRNA were added in combination with RNase H (NEB) to NE. In sham conditions enzyme storage buffer or H2O was used. The targeted digestion of snRNA was performed as described previously (Black et al., 1985, Wongpalee et al., 2016). The treated 20 μl NE aliquots were either used directly or stored in -80°C.

### Complex assembly

Prespliceosomal complexes (10 μl) were assembled on 2.5 nM of [^32^P] labelled pre-mRNA transcript with 50% (v/v) Hela nuclear extract, based on the standard condition used in the *in vitro* splicing reaction. Deviations from the standard conditions are indicated in the figure legends. Reactions were incubated at 30°C for 10 min or as indicated. After complex formation, an additional 10 min incubation was performed with heparin added to the final concentration of 0.5 mg/ml. Similar to the band shift assay, the reactions were chilled on ice, to which 2 μl 50% (v/v) glycerol was added. Complexes were loaded on a pre-run of native PAGE gel, 4%, 80:1, acrylamide:bisacrylamide ratio, using 50 mM Tris-glycine pH 8.8 running buffer, running 160V at RT for 5 hour. Gels were dried on a filter paper, autoradiography was performed as described above.

### Protein-RNA UV crosslinking and Immune-precipitation

20 μl prespliceosomal complexes subjected to UV crosslinking were assembled on 2 nM [^32^P] labeled RNA transcript without PVA, in otherwise identical fashion as those aim to be resolved on an native gel. After complex assembly and incubation with heparin, the reactions were radiated with 2×960mJ 240nm UV-C light. Non-crosslinked RNA were digested by 8 μg RNase A and 0.024U RNase T1 followed by an incubation at 37°C for 12 mins. For immunoprecipitation, RNase treated sample was incubated with 90 μl NETS buffer (10 mM Tris-HCl, 100 mM NaCl, 10 mM EDTA, 0.2% SDS (w/v) and pH 7.4) and 5 μl of antibody or pre-immune serum. After 1 hour incubation at 4 oC, pull-down was performed with 100 μl pre-blocked (NETS buffer with 4 mg/ml BSA (NEB) and 2 mg/ml tRNA (SigmaAldrich)) 0.2% protein G slurry (Cytiva). Following a further 1 hour incubation at 4°C, protein enriched on the beads was washed (3×NETS buffer via centrifugation at 1000*g*, 1 min) and finally released by 30 μl of reducing Laemmli loading buffer. Protein-RNA crosslinks were resolved on 15% SDS-PAGE gels and visualized by autoradiography.

### Psoralen RNA-RNA crosslinking and snRNA identification

2.5 fmole prespliceosomal complexes (10 μl) was assembled on [^32^P] labeled RNA transcript as described above. Psoralen-AMT (SigmaAldrich) (1 μl) was added to the final concentration of 25 μl/ml, heparin omitted. After a further 10 min incubation and complexes were radiated with UV-A light for 20 min, both steps performed on ice. The total RNA content was harvested by standard PK digestion followed by ethanol precipitation with Glycoblue co-precipitant (Thermofisher Scientific). The precipitated RNA pellets were resuspended in H_2_O. Targeted RNase H (NEB) digestions were performed according to manufacturer’s protocol. Three DNA oligonucleotides were used to verify the substrate RNA crosslinking to snRNA, complementary DNA oligonucleotide to GAPDH mRNA was used as an negative control (Table S.4). The digested RNA was purified via standard phenol extraction procedure, ethanol precipitated and analyzed on a 4% denaturing Urea-PAGE acrylamide gel. The crosslinking products and sensitivity to complementary DNA oligonucleotides were determined by autoradiography with a Phospho-imaging screen and imaged with a Typhoon FLA9000 imager.

### Trans-splicing

Transcription templates for AML_E1 and TM4_40exU1 pre-mRNA were generated by oligonucleotide synthesis and cloned into pGEM-4Z vector (Table S.5). All pre-mRNA substrates used in the trans-splicing assay were *in vitro* transcribed with T7 RNA polymerase (Table S.11), 80% capped with m7G cap analog (NEB), and treated with DNase turbo (ThermoFisher Scientific, 37°C, 30 min). After column purification with RNA monach kit (NEB), the concentration of RNA transcripts were determined by UV absorbance. *Trans*-splicing reaction condition is similar to that of *cis*-splicing condition described previously except splicing was performed concurrently with 5 nM of regulated (TM3 or TM23) and 50 nM constitutive (AML_E1 or TM4_40ex) RNA substrates, at 3.6 mM MgCl_2_ and 2 mM ATP. After incubation, spliced products were phenol extracted from the PK digestion, ethanol co-precipitated with 20 μg of glyco-blue (ThermoFisher Scientific). 10 μl RNA product dissolved in water was pre-incubated (65°C, 5 min) with 20 picomole RT primer (Table S.6) and dNTP, cooled on ice, before reverse transcribed with SuperScript II Reverse transcriptase (Invitrogen) according to the manufacture’s instruction.

10% of the RT reaction was used as the template in 25 μl PCR reactions containing 1.25U of JumpStart Taq Polymerase (Sigma D9307), 1×PCR buffer (Sigma P2192), 400 nM of primers (Table S.6) and 0.2 mM dNTP. The reactions were heated (94 oC, 3 min) before 32 amplification cycles (94°C for 30 s, 60 °C for 30 s, and 72°C for 60 s,) and a final extension (72°C, 60 s). PCR products were subsequently resolved on the QIAxel advanced system (QIAGEN) using a DNA screening capillary electrophoresis cartridge.

### Affinity enrichment of prespliceosomal proteome

RNA assisted pull-down was adapted from (Jurica et al., 2002) and performed under 8 conditions, each contains three technical repeats as detailed in Fig.S12B. Before purification, *in vitro* transcribed TM3-MS2 RNA was heated at 80°C for 2 min and refolded at RT for 5 min in buffer RB (20 mM HEPES, 0.2 mM EDTA and pH 7.9). MS2-MBP protein (MS2 bacteriophage coat protein-maltose binding protein fusion) binding was performed in RT of 15 min at a protein to RNA ratio of 20:1. After thawing, a 17000*g* 5 min centrifugation of the Hela nuclear extract was performed to remove aggregation. A 400 μl binding reaction was constituted, in condition similar to that of the *in vitro* splicing reaction, with 200 μl of clarified NE, ±5 nM RNA, ±1.5 μM recombinant RBPMS, ±0.5 mM ATP, ±20 mM creatinine phosphate, 3 mM MgCl_2_, 70 mM KCl, PVA omitted and pH 7.9. After incubation for 20 min at 30 °C, the binding reaction was added to 200 μl of 5% (v/v) amylose beads (NEB) that has been blocked overnight with 1 mg/ml BSA and 0.5 mg/ml tRNA in 20 mM HEPES, 70 mM KCl and pH 7.9. 4× washes were conducted following 1 hr incubation at 4°C, using WB-50 buffer (20 mM HEPES, 50 mM KCl and 0.5 mM DTT pH 7.9). To elute, the volume was opted, 10× of the beads volume, using WB-50 buffer supplemented with 40 mM maltose (SigmaAlrich).

### Affinity enrichment of RBPMS proteome

Purified StrepII-His_6_-RBPMS-A protein was exchanged into buffer BS using a Zeba spin desalting column (7K MWCO, ThermoFisher Scientific). Pull-down assay were assembled as 166 μl reactions containing 60% Hela NE, 2 μM StrepII-His_6_-RBPMS, 2.2 mM MgCl_2_ and 2.6% PVA. Where applicable, Hela NE was pre-treated with 5U/ml Benzonase (Milipore) at 30°C for 15 min and clarified by centrifugation (17500*g*, 5 min) prior to the addition of RBPMS protein. The pull-reaction were incubate at 30°C for 15 min, added to 200 μl of 2.5% (v/v) of MagStrep “type3” XT beads (IBA Lifescience) that has been blocked overnight with 1 mg/ml BSA (NEB) and 0.5 mg/ml tRNA (SigmaAlrich) in buffer WB-150 (20 mM HEPES, 150 mM KCl and 0.5 mM DTT pH 7.9), and further incubated at 4°C for 1hr. After removal of the flow-through, beads were washed six times with buffer WB-150 (6 × 1ml). The elution was carried out at RT with shaking for 30 min by adding 45 of elution buffer (100 mM Tris, 150 mM NaCl, 1 mM EDTA, 50 mM Biotin, pH 8).

### Mass spectrometry

For RNA assisted pull-down, LC MS/MS was used to identify and quantify proteins recovered from RNA assisted pull-down. Elute was trichoroacetic acid (TCA) precipitated, re-dissolved in reducing Laemmli loading, separated by SDS-PAGE and visualized with silver staining (Fig.S12C). The serial gel slices were excised and digested *in situ* with trypsin. The extracted tryptic peptides were analyzed using Q-Exactive mass spectrometer. Raw data were processed using ProteomeDiscoverer V2.3 (Thermo Fisher Scientific). Protein identification was conducted by searching Human database downloaded in 2020, UniPort, using Mascot algorithm. This generated a list of 1081 entries containing common contaminant proteins (human keratins, heat-shock proteins and BSA), which were identified and removed from downstream analysis. The data obtained from ProteomeDiscoverer was abundance data at peptide level. Data was processed with R package and filtered to remove entries that only identified in 1 out of 3 replicates of at least one conditions. The resulting 978 entries (supplementary file RNA_MS_A) was background corrected and normalized by variance stabilizing transformations. Inspection of the list revealed repetitive interpretation due to isoforms of the same protein and searching multiple databases. We collapsed the repetitive isoform entries of the same protein, shortlisted 178 unique identifications for further analysis (supplementary file RNA_MS_B). Low intensity missing values were biased to no RNA background samples and no RBPMS added conditions. To conduct the differential expression (DEP) analysis, missing Total Precursor Intensity was imputed using random draws from a Gaussian distribution centered around a minimal value, q-th quantile = 0.01. We used R package <u>L</u>imma to test the significant changes between background subtracted groups as tableted in Fig.S12B. The fold-changes were estimated by the Bayes method, while the adjusted *P-value* were corrected by the Benjamini-Hochberg method.

For AP-MS experiment, the sample preparation, peptide identification and raw data processing is identical to that described above. Initial proteomic data processing was carried out in Scaffold (Searle, 2010) (supplementary file AP_MS_TSC_raw). Entries detected in the lack of nuclear extract condition (Data file (supplementary file AP_MS_TSC_raw, samples BL1-3) were excluded from analysis. To further enriched the list of significantly recovered proteins, total spectrum count of grouped technical repeats were compared using unpaired student’s T-test. FL vs negative control produced 131 significant interactors (P < 0.05) while vs ΔC20 generated 133 significant interactors (supplementary file AP_MS_SL). ΔC20 vs negative control produced 48 significant interactors. GO analysis was performed on STRING with the following parameters adjusted: interaction sources set to experiments and databases only, minimum required interaction score set to medium confidence (0.400).

### Glycerol gradient ultracentrifugal sedimentation

For analysis of RBPMS-A associated protein complexes in Hela NE, 13 ml 10% – 60% w/v glycerol, 0% - 0.15% glutaraldehyde gradient (20 mM HEPES, 100 mM KCl, 2.2 mM MgCl2, 0.5 mM DTT, 0.2 mM EDTA, pH 7.9) were set up in 14 × 95 mm tubes (Beckman Coulter). 166 μl reactions were assembled as described previously for Benzonase-treated affinity pull-down assay, but instead of adding 2 μM StrepII-His6-RBPMS, a mixture of 1 μM StrepII-His6-RBPMS-A (Repeat 1) or tag free RBPMS-A (Repeat 2-3) and 1 μM His6-RBPMS-A conjugated to Alexa-647 dye was used. Samples were loaded onto the gradient and subjected to ultracentrifugation in a SW40 Ti rotor (Beckman Coulter) at 32,000 RPM for 13 hr at 4 °C. Gradients were fractionated into 24 × 0.5 ml fractions. The sedimentation coefficient was deduced by analysing E. coli lysate (Rivera et al., 2015), the 30S or 50S ribosomal subunit fractions were identified by determination absorption at 260 nm. 50 μl of each fraction was diluted with 2 × transfer buffer (50 mM Tris/glycine, 16% v/v methanol, pH 8.8) and dot blotted onto an immobilon-FL PVDF membrane (Milipore). Immuno-detection was carried out using the antibody and dilution tabulated (Supplementary S.10). Chemiluminescence was developed using Clarity Max ECL substrate (Bio-rad) and imaged via Bio-rad ChemiDoc MP imager. Imaging processing was conducted using ImageStudioLite package (LI-COR).

### PAC-1 cell culture and RNAi gene silencing

Rat pulmonary artery smooth muscle PAC-1 were cultured following standard procedures to maintain the differentiated state (Llorian et al., 2016). siRNA mediated knockdown was performed using reverse transfection. Briefly, 60 - 90 pmol siRNA and Lipofectamine RNAiMAX reagent (ThermoFisher Scientific) were diluted by Opti-MEM without serum then mixed and incubated at RT 20 min. Diluted 2.5 × 105 differentiated PAC-1 cells per well were added to the RNAi duplex - Lipofectamine RNAiMAX complexes. An siRNA of scrambled sequence “C2” was used as control in every set of knockdown experiments. All siRNA sequences were tabulated in Table S.8. Each condition was repeated in ×6 format, to allow triplicates of RT-PCR analysis and sufficient material for verification of the knockdown at protein level. In the two-hit experiment, the cells were treated again with the same reagent and procedure after 24 hr. Cells were harvested 48 hr after the terminal siRNA treatment.

### RT-PCR and RT-qPCR

To verify the silencing of target gene, cDNA was prepared using 1 μg of total RNA, oligo(dT) (Merck) and superscript II (Invitrogen) based on the instruction given by the manufacturers. RT-qPCR was performed as in (Nakagaki-Silva et al., 2019), but using gene specific primers (Supplementary Table S.9). Two housekeeping genes were included in each analysis (CANX and Rpl32), their geometric means were used to normalize the relative expression values. Expression values were acquired from three biological repeats.

To examine the usage of differentiation specific exon usage, PCRs with 50 ng of cDNA were performed using the oligonucleotide primers listed in supplementary Table S.9. The PCR products were resolved in a QIAxcel system as described previously. The visualization and quantification of percentage of splicing in (PSI) values were conducted using QIAxcel ScreenGel software. PSI values are expressed as mean (%) standard deviation (SD). Statistical significance was examined with unpaired Student’s T-test.

**Table S.1.**
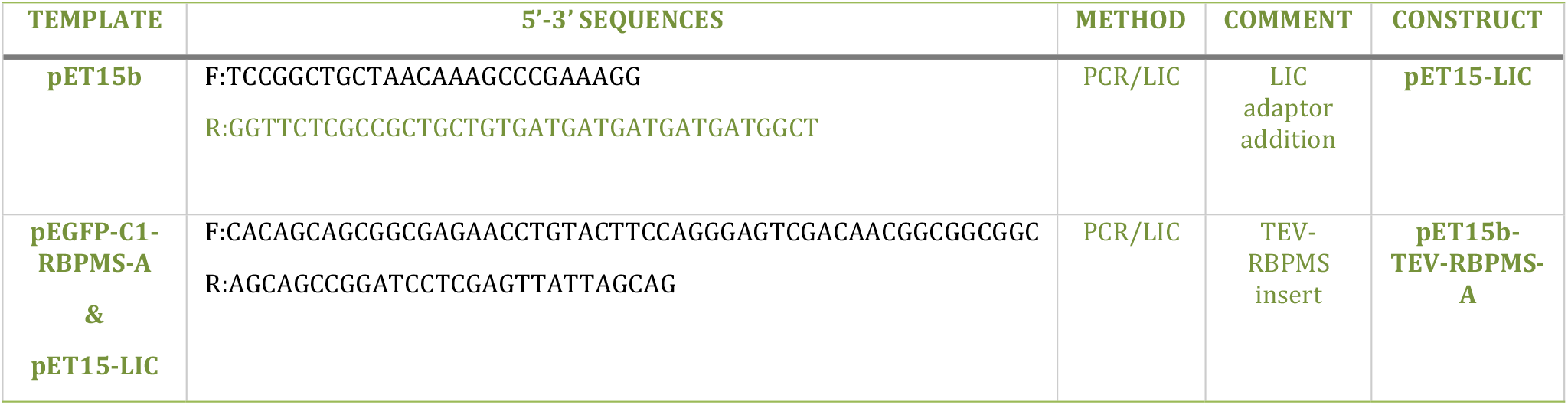
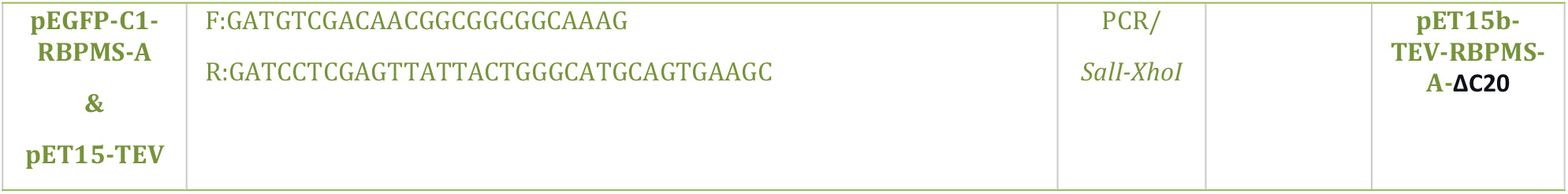
Primers used in cloning of RBPMS constructs with TEV cleavable His6 tag. LIC, ligation independent cloning; F, Forward ; R, Reverse.

**Table S.2.**
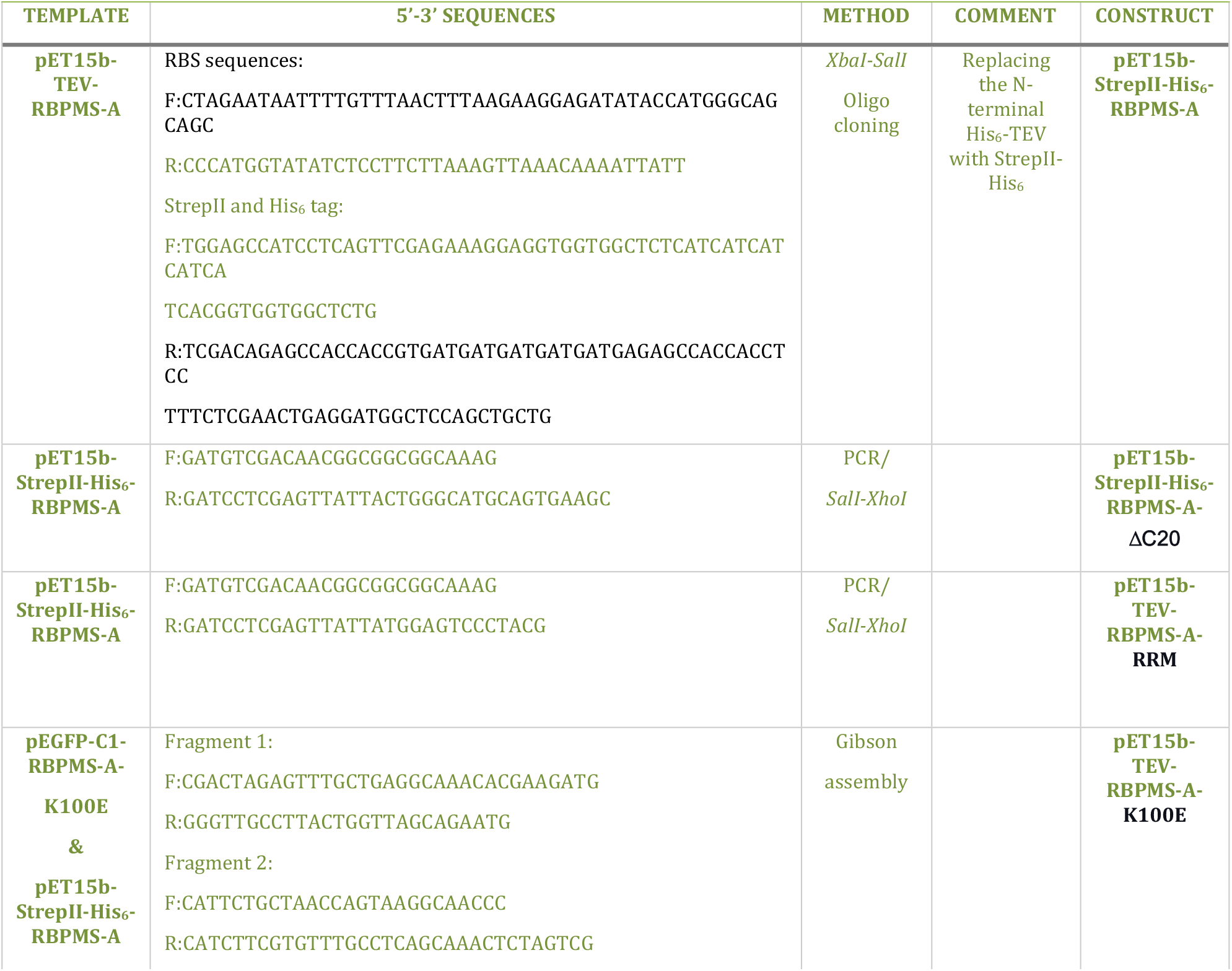
Primers used in cloning of RBPMS constructs with N-terminal StrepII and His_6_ tags. RBS, ribosome binding sequence ; F, Forward ; R, Reverse.

**Table S.3.**
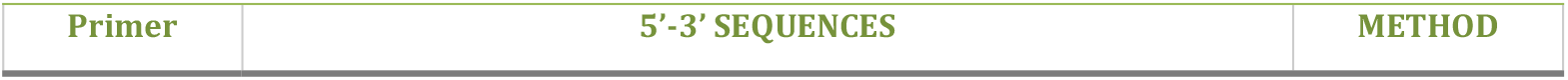
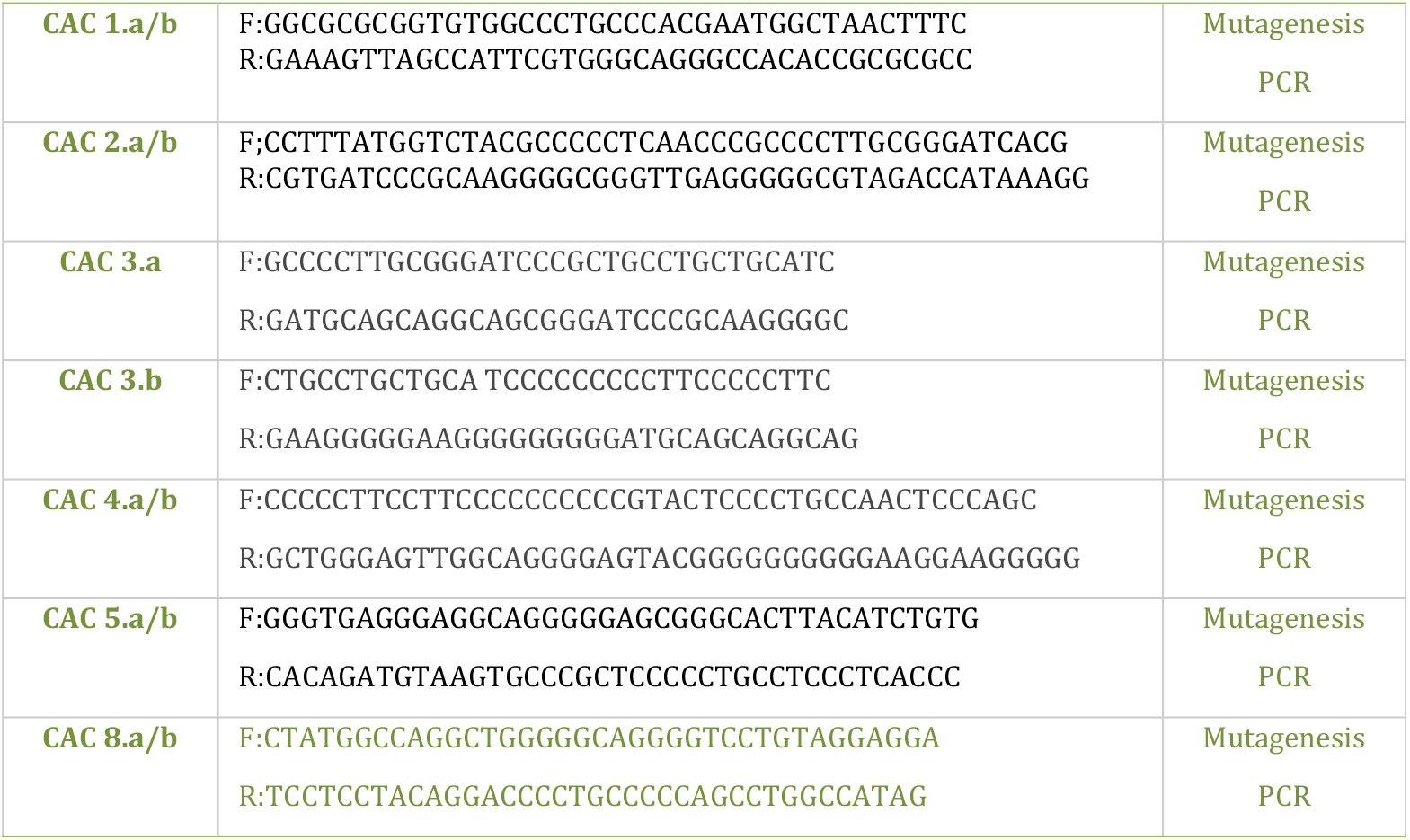
Primers used in the depletion of tandem CAC sites; F, Forward primer; R, Reverse primer.

**Table S.4.**
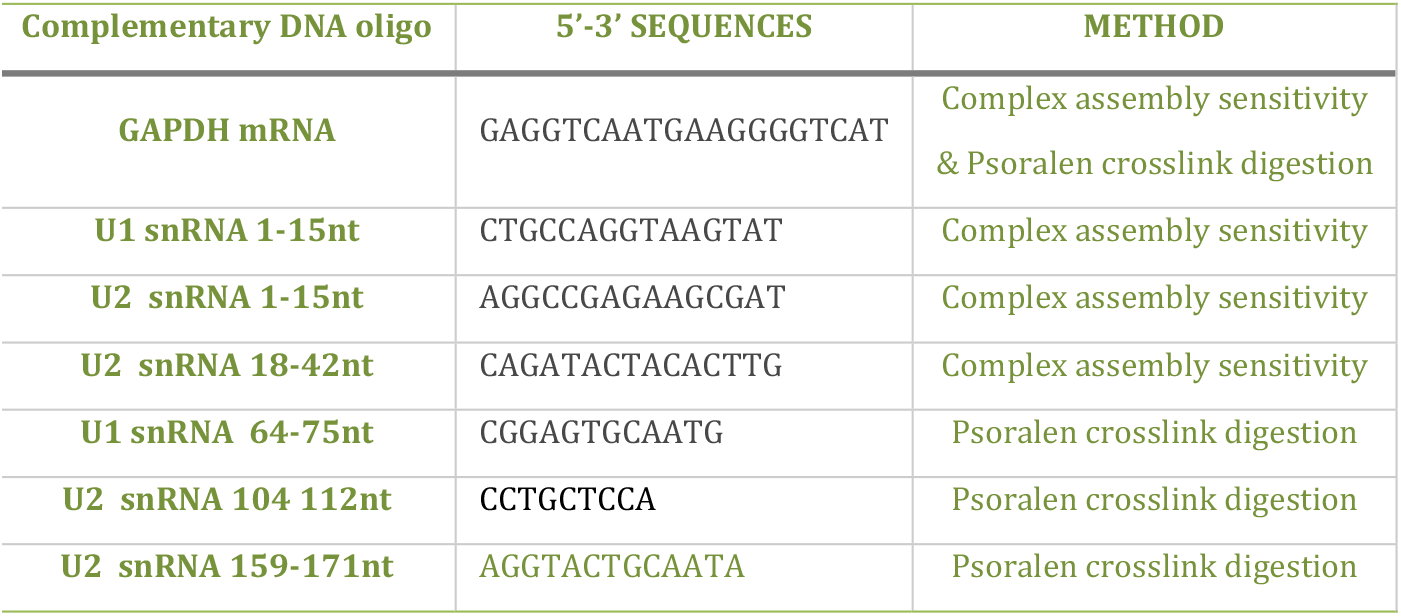
List of complementary DNA oligonucleotide used in combination with RNase H.

**Table S.5.**
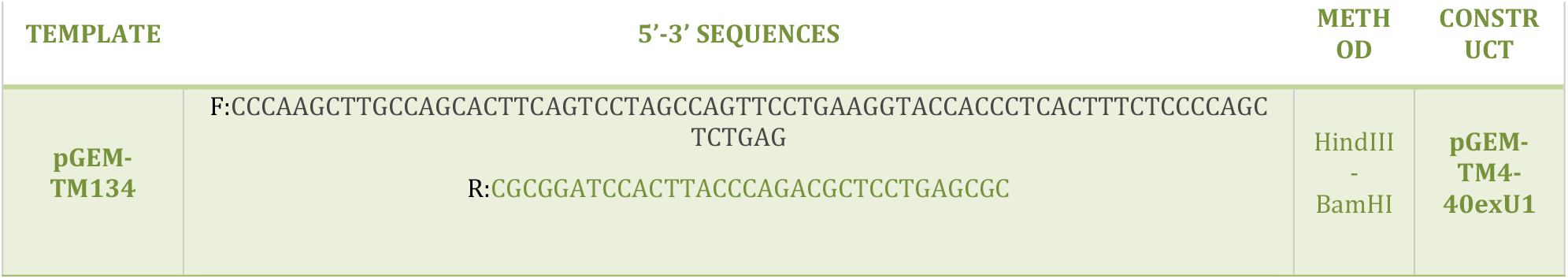
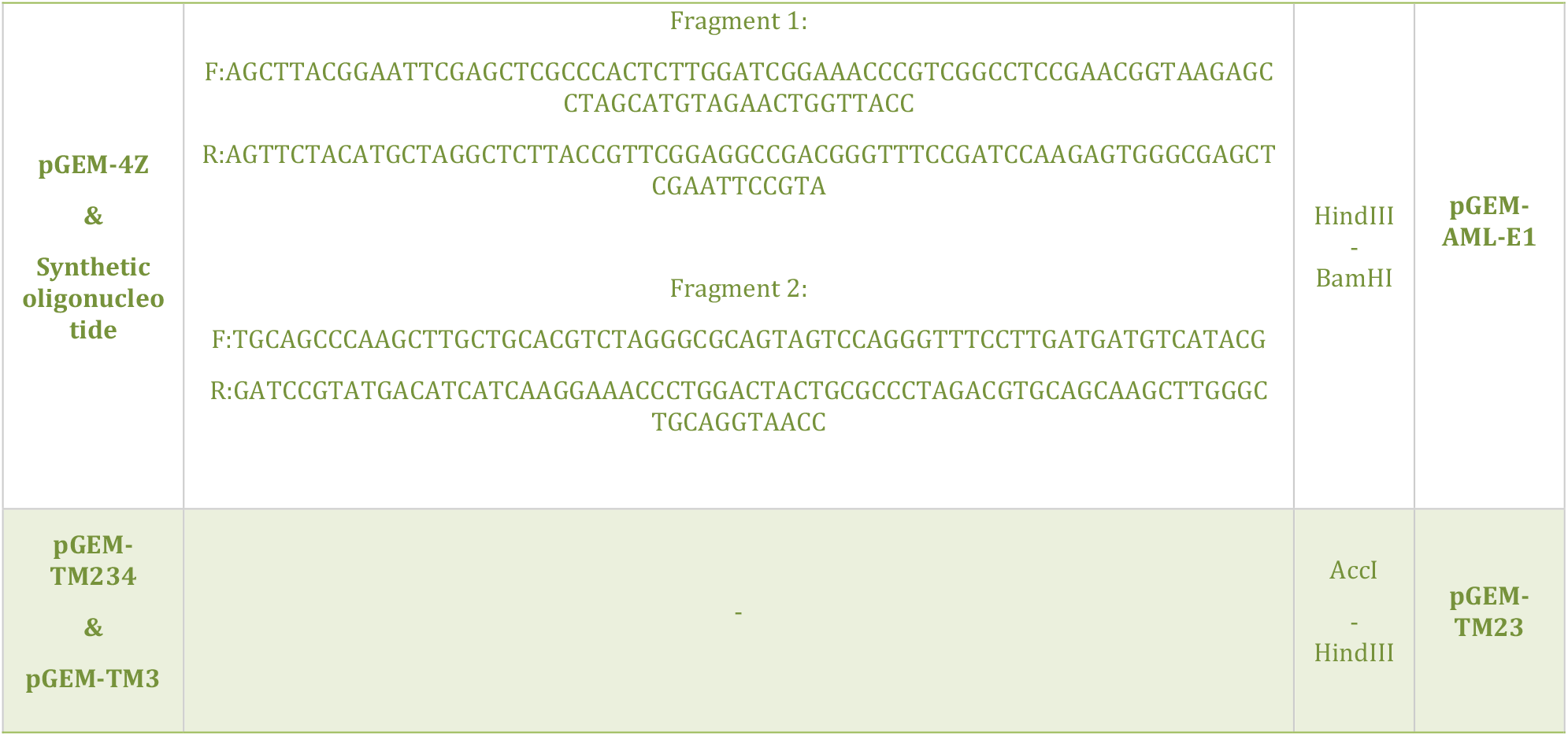
Cloning strategies for splicing substrates; F, Forward primer ; R, Reverse primer.

**Table S.6.**
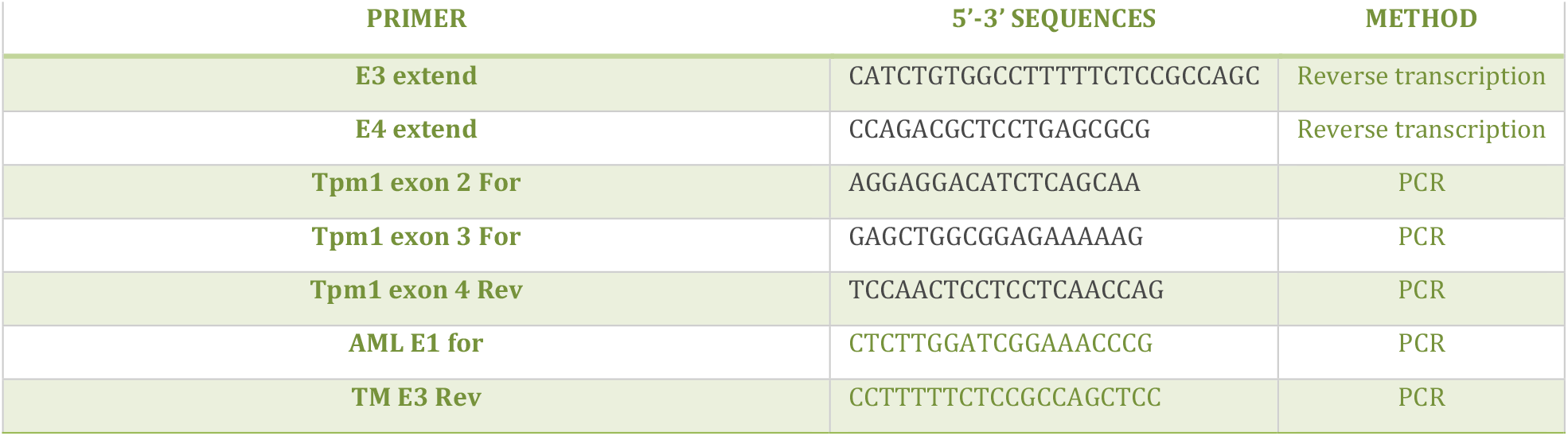
List of primers used in *trans* splicing product detection.

**Table S.7.** Linearized template plasmid for *in vitro* transcription.

| Experiment | Plasmid | Restriction enzyme | Capping % |
| --- | --- | --- | --- |
| Cis-splicing | pGEM-TM134 | XbaI | 50 |
|  | pGEM-TM234 | BamHI | 50 |
| Complex assembly &<br>*Trans*-splicing<br>UV crosslinking | pGEM-TM3 | EcoRI | 80 |
|  | pGEM-TM23 | EcoRI | 80 |
| RNA affinity pull-down | pGEM-TM3-MS2 | EcoRI | 80 |
| *Trans*-splicing | pGEM-TM4-40exU1 | BamHI | 80 |
|  | pGEM-AML-E1 | BamHI | 80 |
| Band shift | pGEM-P3CAC | XbaI | 0 |
|  | pGEM-SCAC | EcoRI | 0 |

**Table S.8.**
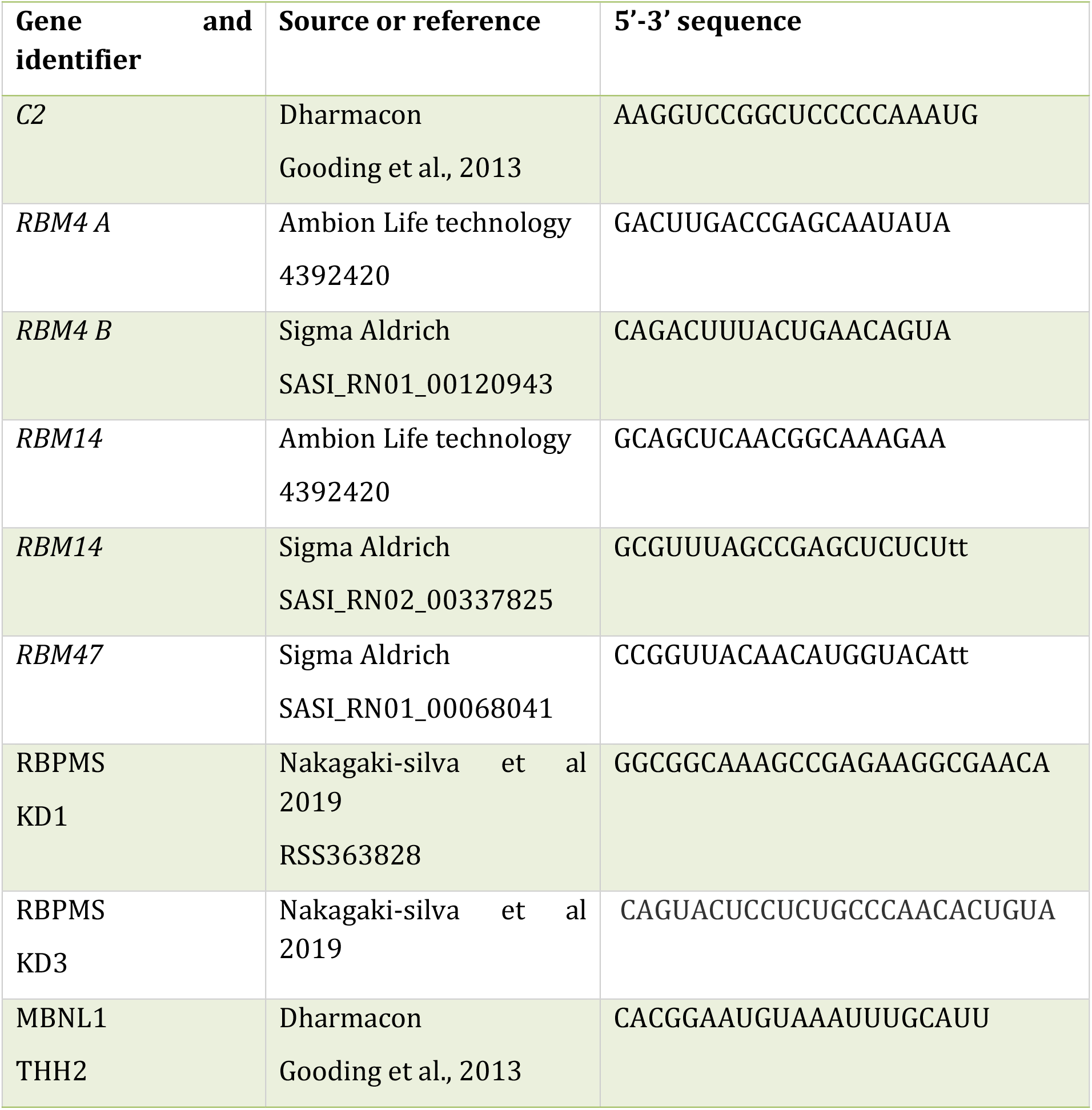
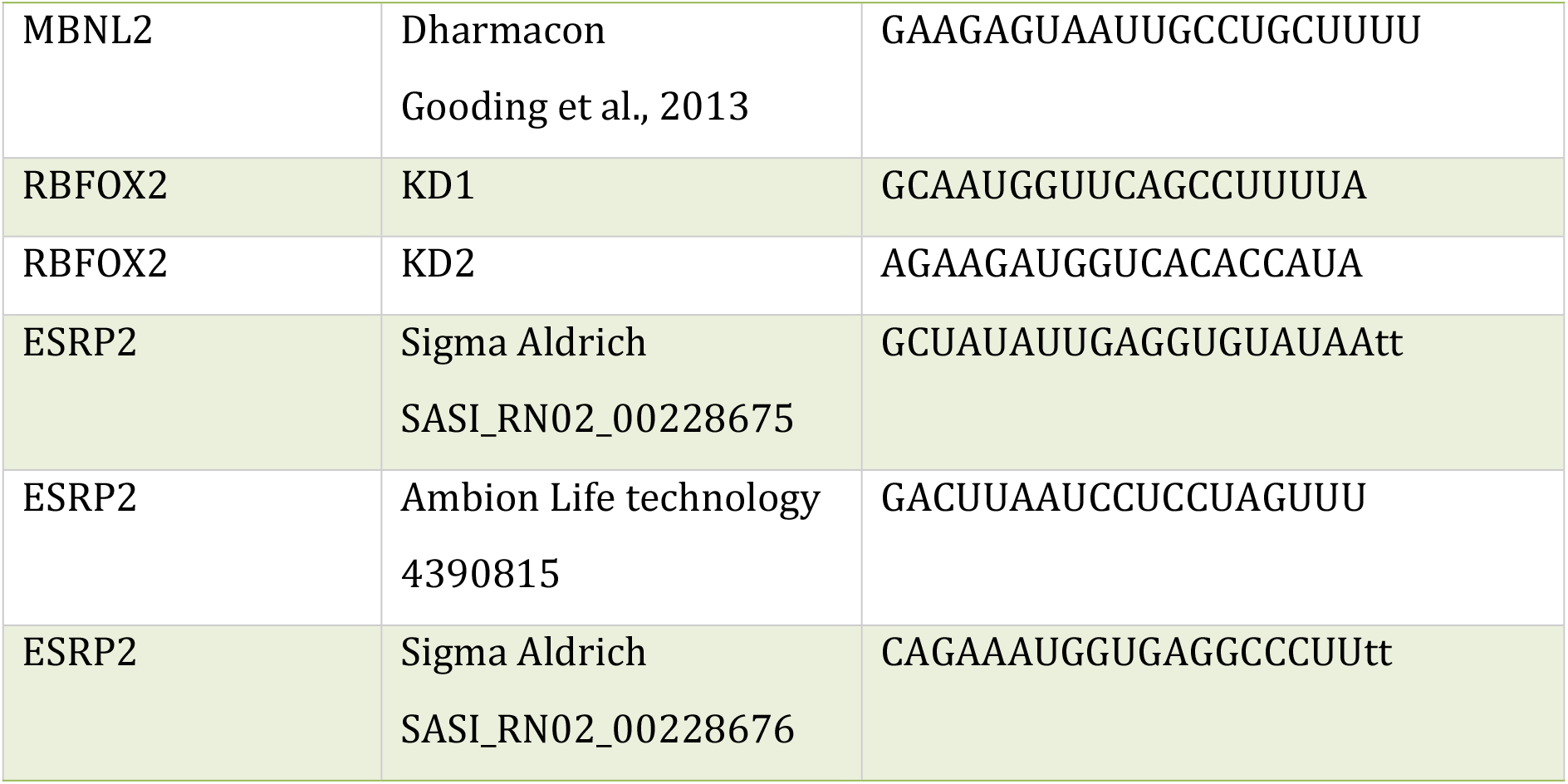
List of siRNA.

**Table S.9.**
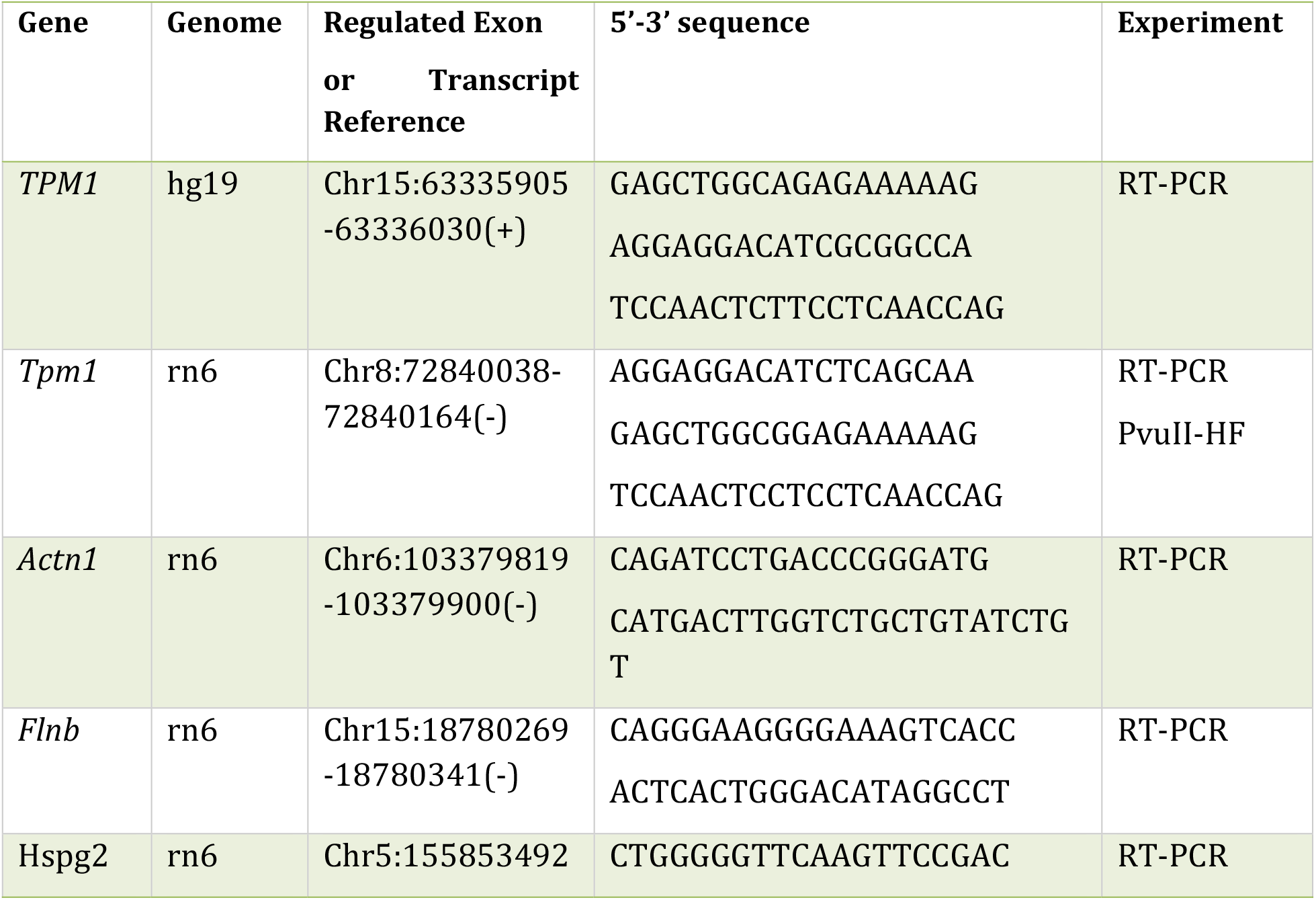
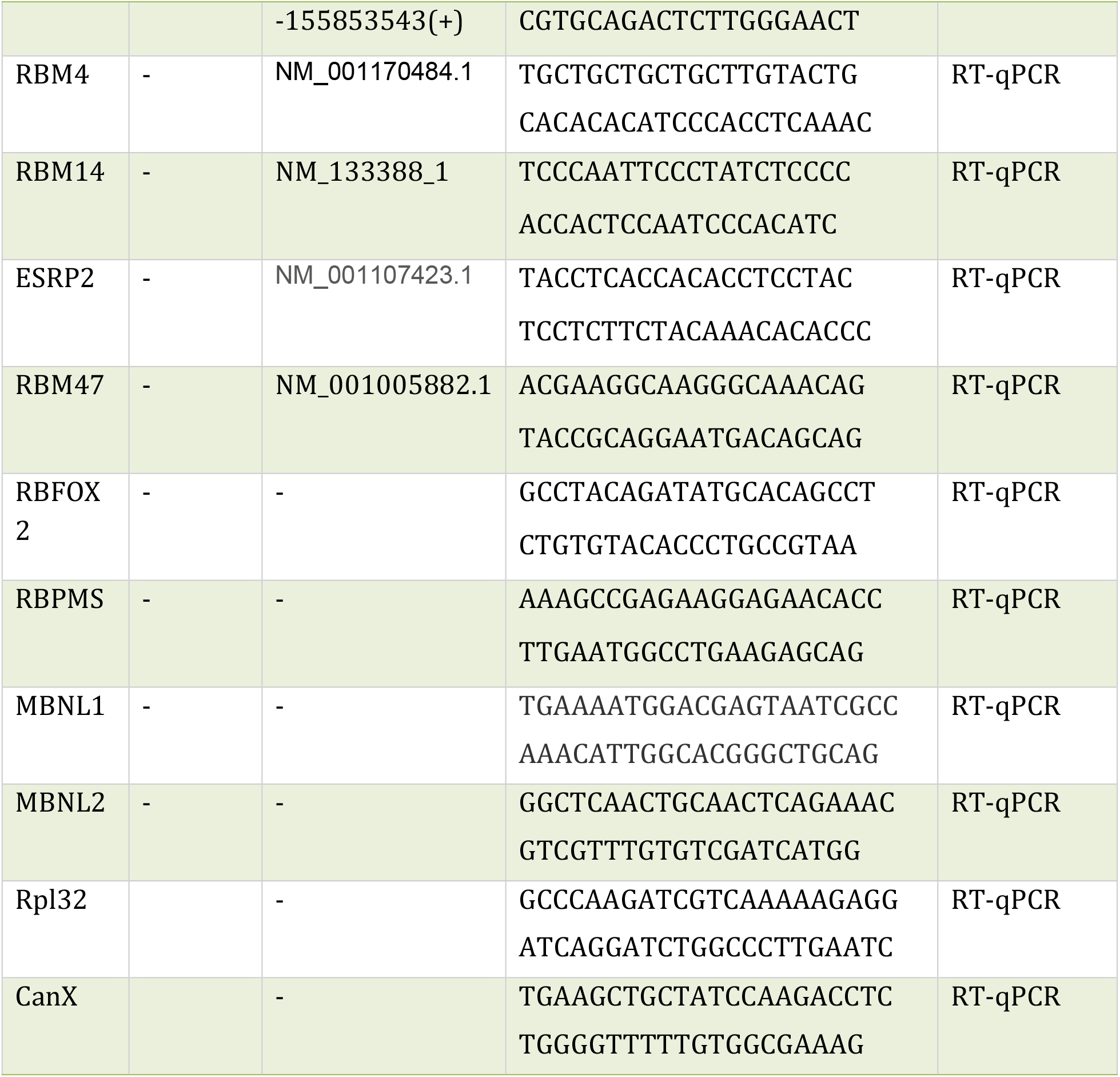
List of PCR primers for knockdown validation.

**Table S.10.**
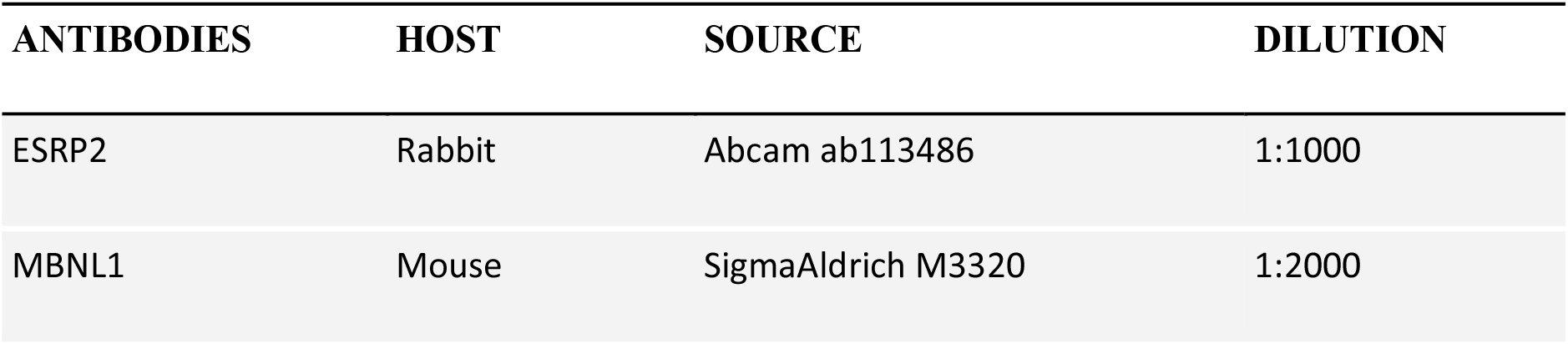
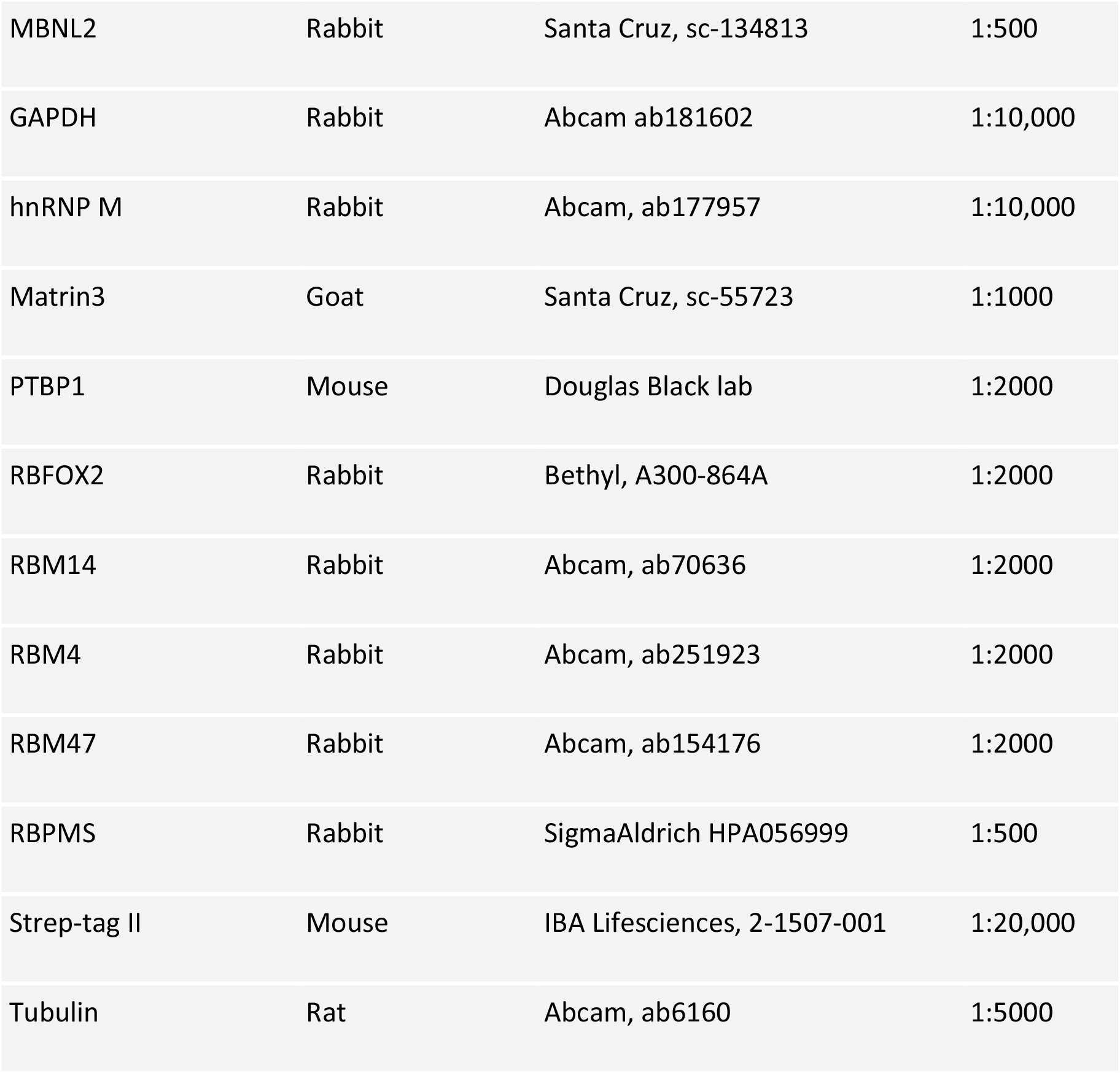
List of antibody.

**Table S.11.** In vitro transcription.

|  | Band shift | Cis Splicing substrates | Complex assembly & Psoralen crosslinking | UV Crosslinking | Trans splicing Substrates & RNA affinity pull-down |
| --- | --- | --- | --- | --- | --- |
| Capping % (Radio-specificity) | 0 (low) | 50(low) | 80(low) | 80(high) | 80(no) |
| Linearised RNA 1 mg/ml (µl) | 1 | 1 | 1 | 1 | 1 |
| 1 µl of 5 mM A,U,GTP (x mM CTP) | 5(2) | 5(2) | 5(3) | 5(0.4) | 10 µl of 10 mM A,U,CTP (2 mM GTP) |
| Ribolock RNase Inhibitor (μl) | 0.4 | 0.4 | 0.4 | 0.4 | 1 |
| 5x transcription buffer (μl) | 2 | 2 | 2 | 2 | 10 |
| [32P] CTP (μl) | 1 | 1 | 1 | 2 | - |
| 10 mM Cap analogue (μl) | - | 0.5 | 0.8 | 0.8 | 8 |
| T7 RNA polymerase (μl) | 1 | 1 | 1 | 1 | 1.5 |
| H2O | 3.6 | 3.1 | 2.8 | 1.8 | 18.5 |

## Acknowledgements & Funding

For the purpose of open access, the authors have applied a Creative Commons Attribution (CC BY) license to any Author Accepted Manuscript version arising from this submission. We thank Ian Eperon and Alex Borodavka for helpful and insightful comments on the manuscript. This work was funded by a Wellcome Trust Investigator award (209368/Z/17/Z) to CWJS. EENS was supported by a studentship from the Conselho Nacional de Desenvolvimento Científico e Tecnológico (206813/2014-7) and YH by a Chinese Scholarship Council Cambridge International Scholarship.

## Conflicts of Interest

The authors declare that they have no conflict of interest

